# Somatic mutant selection is altered by prior *NOTCH1* mutation in aging esophageal epithelium

**DOI:** 10.1101/2024.09.26.615123

**Authors:** Charlotte King, Emilie Abby, Joanna C. Fowler, Kasumi Murai, Roshan K. Sood, Irina Abnizova, Swee Hoe Ong, Salomé Brunon, Faye Lynch-Williams, Michael W. J. Hall, Benjamin A. Hall, Philip H. Jones

## Abstract

The ageing mid esophagus is progressively colonized by somatic mutations, including oncogenic *TP53* and antioncogenic *NOTCH1* mutants. Most esophageal squamous carcinomas are *TP53* mutant, *NOTCH1* wild type. By middle age, much of the esophagus is *NOTCH1* mutant. We hypothesised that prior *NOTCH1* mutation may alter subsequent mutant selection. Here we show that selection of mutant *NOTCH2* was increased and mutant *TP53* decreased within *NOTCH1* mutant epithelium. Potential mechanisms include spatial competition between adjacent clones and/or epistasis altering the fitness *NOTCH1* double mutants. We investigated these possibilities in transgenic mice. Spontaneous *Notch2* mutant clones emerged in aged *Notch1* null, but not wild type, esophagus, consistent with epistasis increasing *Notch1/Notch2* double mutant fitness. However, there was no evidence of epistasis between *Notch1* and *Trp53,* as *Notch1* mutation alone increased fitness. In consequence, over time, *Trp53* mutant *Notch1* wild type clones were outcompeted by expanding *Notch1 and Trp53/Notch1* mutants. *NOTCH1* mutant takeover of the aging esophagus alters the trajectory of somatic evolution, decreasing selection of *TP53* mutants and potentially reducing cancer risk.

## Introduction

Somatic evolution in normal epithelia occurs as progenitor cells accumulate mutations due to cell intrinsic processes and environmental exposures^1, 2, 3^. A subset of mutant genes alters progenitor cell behavior to drive clonal expansion. In turn, this leads to selection as colliding clones compete for space within the progenitor cell niche. The fittest clones survive to acquire further mutations while less fit mutants are lost. This process is well illustrated in the aging mid esophagus, which develops into a patchwork of mutations from which esophageal squamous cell carcinomas (ESCC) may emerge^3, 4, 5, 6, 7, 8^.

The most prevalent mutant gene in the aged esophagus is *NOT*C*H1*, followed by *TP53*. However, despite most of the older esophagus being mutant for *NOTCH1*, only 15% of ESCC carry *NOTCH1* mutations (**Supplementary Fig. 1a**)^3, 7, 8^. This implies that *NOTCH1* mutation may protect against malignant transformation, a hypothesis consistent with observations in carcinogenesis models in mice^7, 9^. Substantially less epithelium is mutant for *TP53* than for *NOTCH1*. However, most premalignant lesions and over 90% of ESCC harbor nonsynonymous mutation and/or loss of heterozygosity (LOH) of *TP53*^10, 11^. Consistently, mouse models indicate *Trp53* mutation contributes to multiple processes in esophageal squamous carcinogenesis^5^. Most ESCC develop from *TP53* mutant, *NOTCH1* wild type clones, so processes that limit the size of this population may reduce cancer risk (**Fig. 1a**)^5, 10, 11^.

**Figure 1:**
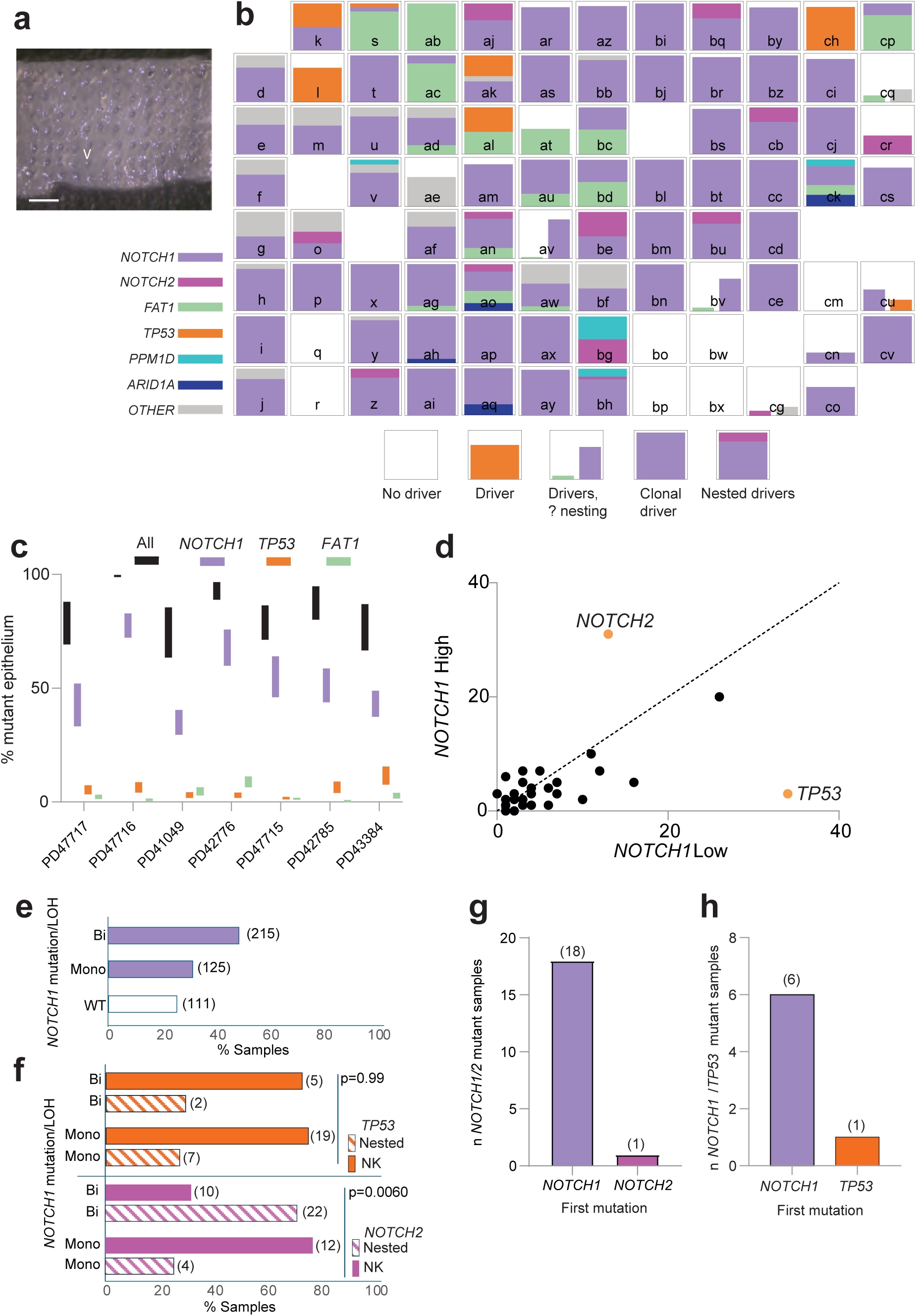
Mutations in 0.05mm^2^ samples. **a** 0.05 mm^2^ sample collection. A regularly spaced array of samples was collected human esophageal epithelium using a circular 250mm diameter punch (arrow) from a 70-year-old male (PD42776) scale bar: 1 mm. **b** Targeted sequencing of tissue shown in **a** for 324 cancer associated genes. Bar charts show proportion of sample mutant for positively selected mutants shown. Position of bar chart corresponds to the spatial location of the sample. Letters indicate sample identity. Bar height indicates proportion of mutant tissue derived from variant allele frequency (VAF). Stacked bars indicate nested mutations (the lowest bar is clonal). Side-by-side bars indicate that nesting is unknown. Color indicates mutant gene. ‘other’ indicates mutation in another positively selected mutant gene, **c** Proportion of mutant tissue in all samples from each donor derived from variant allele frequency. Three most prevalent mutant genes are shown, ‘All’ is all positively selected nonsynonymous mutations. Bars indicate range of possible values due to uncertainties over copy number and nesting. Lower limit shows the result of biallelic mutation of the gene in all mutant cells with copy neutral loss of heterozygosity or compound heterozygous mutations. Upper limit corresponds to one mutant allele per cell at a diploid locus. **d** Mutant selection in 0.05 mm^2^ samples with different levels of *NOTCH1* mutation. Samples were sorted by the summed VAF of all *NOTCH1* mutants they carried into mutant *NOTCH1*High (*NOTCH1* summed VAF >0.4) and *NOTCH1*Low (*NOTCH1* summed VAF <0.4). Number of protein-altering mutations is plotted for each sequenced gene with 5 or more mutations in the *NOTCH1High* or *NOTCH1Low* groups, orange indicates a significant difference. p values, mutant *NOTCH2,* 0.00347, *TP53* <0.0001, 2 tailed Z test. n samples *NOTCH1*High 239, *NOTCH1*Low 212. **e** Percent of 0.05 mm^2^ samples that are *NOTCH1* wild type or carry monoallelic (Mono) or biallelic (Bi) mutation/LOH **f** Nesting of *TP53* and *NOTCH2* mutants within *NOTCH1* mutant clones with monoallelic (Mono) or biallelic (Bi) *NOTCH1* mutation/LOH. Samples containing nested mutants and samples in which nesting cannot be determined (NK) for samples with *NOTCH*2 and *NOTCH1* or *TP53* and *NOTCH1* mutants. Number of samples in brackets, Fisher’s exact test. **g**, **h** order of mutations in nested double mutant samples in which this could be determined. **g** *NOTCH1/NOTCH2*, **h** *NOTCH1/TP53*, number of samples in brackets.

Potential mechanisms underpinning selection somatic evolution in the esophagus include spatial competition between adjacent clones and epistasis as mutant cells accumulate additional mutations^5, 9, 12, 13^. The extensive colonization of the esophagus by *NOTCH1* mutants by late middle age raises the possibility that the selection of mutants over subsequent decades may diverge in *NOTCH1* mutant and wild type epithelium. Here we investigate this hypothesis by spatial mapping of clones in aged human esophagus and exploring selective mechanisms in transgenic mouse models.

## Results

### Mapping mutations in aged human esophagus

Existing mutational data from human esophagus was too sparse for reliable detection of effects such as epistasis^3, 8, 14^. We therefore sequenced normal mid esophagus from 22 transplant donors with no history of cancer or esophageal disease aged up to 79 years, combining the results with nine previously reported donors sequenced using the same methods (**Supplementary Fig.1b, Supplementary Table 1**)^8^. Analysis of germline single-nucleotide polymorphisms altering the risk of developing ESCC showed the donors were genetically representative of the UK population (**Methods**, **Supplementary Table 2**)^15, 16, 17, 18, 19^. In total of 451 0.05 mm^2^ and 2405 2 mm^2^ samples in spatially mapped arrays were sequenced (**Fig. 1b, methods).**

We began by determining whether the mutational features in our samples of esophagus were consistent with those described in previous studies. To measure mutational burden and identify mutational signatures we used Nanorate sequencing (NanoSeq), a modification of duplex sequencing that detects sequence variants across 30% of the genome with high accuracy ^1^. This showed that mutational burden increased linearly with age, with a higher burden in ever smokers (**Supplementary Fig. 1c, Supplementary Table 3**). Three age correlated single base mutational signatures (SBS1, 5 and 40) were detected, consistent with previous studies^1, 3, 8, 20^ (**Supplementary Fig. 1d**). Notably we did not detect a tobacco related mutational signature in smokers, but this finding agrees with previous studies of normal esophageal epithelium, premalignant lesions and ESCC where tobacco signatures were not found in most smokers^3, 8, 11, 21^.

We also examined mutant selection using targeted sequencing. To identify which mutant genes were under selection, we calculated the ratio of nonsynonymous to synonymous mutations (*d*N/*d*S), adjusted for the mutational spectrum and other factors, for each gene^22,23^. Genes with a dN/dS ratio significantly above 1 (q<0.01, dNdScv) were considered positively selected. Allowing for differences in the panel of sequenced genes, the same genes were selected in the 0.05 mm^2^ and 2 mm^2^ samples. Selected genes were as expected from previous studies with two additional mutant genes, *EEF1A* and *EPHA2* found to be positively selected (**Supplementary Fig. 1e, f, Supplementary Table 4**)^24, 25^. We concluded that the esophageal epithelium sampled here was typical in terms of mutational processes, mutational burden and mutant gene selection.

### Mutational selection is altered by prior *NOTCH1* mutation

To investigate whether prior *NOTCH1* mutation altered mutant selection in the aging esophagus, we first mapped the mutations in 451 spatially mapped 0.05 mm^2^ samples from 7 donors over 60, sequenced for 324 cancer associated genes (**Fig. 1a**). Most (92%) samples contained one or more positively selected mutants, and over half of these were found to be clonal after adjustment for Copy Number Alterations (CNA) (**Fig. 1b, Supplementary Fig. 2a, b Supplementary Tables 4-7)**. Most of the clonal samples carried one or more subclonal mutations (**Fig. 1b**, **Supplementary Fig 2a, b**). The proportion of epithelium occupied by nonsynonymous positively selected mutants approached 100%, with *NOTCH1*, *TP53* and *FAT1* the most frequent mutants by area (**Fig. 1c, Supplementary Table 8**).

As a proxy for clone size, we examined the number of adjacent samples carrying clonal mutations. This ranged widely for commonly mutated genes (**Supplementary Fig. 2c**). As expected, the largest clones carried *NOTCH1* mutants (p=0.0017, Kruskal-Wallis test). The largest mutant clones in each patient were unlikely to form by chance (p<1×10^−6^ for all patients, 2 dimensional permutation test, **Supplementary Fig. 3a-c**). Mutant clones larger than those formed by neutral drift have also been observed in densely mutated esophageal epithelium in mutagenized mice, in which there is strong spatial competition^4^.

We then analyzed the selection of mutant genes in wild type and *NOTCH1* mutant tissue, by comparing the number of nonsynonymous mutants in the 324 sequenced genes in samples that were *NOTCH1* mutant (*NOTCH1*High, i.e. the summed VAF for all *NOTCH1* mutant clones exceeded 0.4, **methods, Supplementary Table 6**) with the remaining *NOTCH1*Low samples. *NOTCH2* mutants were enriched and *TP53* mutants depleted in *NOTCH1*High compared to NOTCH1Low samples (P=0.04, 2 tailed Z test) (**Fig. 1d**). Furthermore, samples containing any *NOTCH1* mutation were more likely to harbor a *NOTCH2* mutation and less likely to carry a *TP53* mutation than *NOTCH1* wild type samples **(Supplementary Fig. 4, Supplementary Table 5).** These results support the hypothesis that prior *NOTCH1* mutation alters the selection of subsequent *NOTCH2* and *TP53* mutants.

Next, we investigated if *NOTCH1* gene dosage impacted mutant selection. There were significantly more *NOTCH1* mutant clones with biallelic compared to monoallelic *NOTCH1* gene disruption, consistent with reports that biallelic loss of *NOTCH1* increases competitive fitness (Chi Square test, p=5.4×10^−5^, **Fig. 1e, Supplementary Table 9**)^7^. Applying pigeonhole analysis to identify nested mutants, we found nested *NOTCH2* mutants were more frequent within *NOTCH1* mutant clones with biallelic compared to monoallelic disruption (Fisher’s exact test, p=0.006, **Fig. 1f, Supplementary Table 9**)^26^. This was not the case for *TP53* mutants (Fisher’s exact test, p=0.99, **Fig. 1f, Supplementary Table 9**).

In some samples it was also possible to infer the order of mutations. *NOTCH1* was the first mutation in 18/19 *NOTCH1/NOTCH2* mutant samples (p=0.0001 by two-tailed binomial test) (**Fig. 1g Supplementary Table 10**). There were only seven *NOTCH1/TP53* double mutant samples, *NOTCH1* mutation occurring first in six of them (p = 0.065 by two-tailed binomial test, **Fig. 1h, Supplementary Table 10**). Thus, *NOTCH2* mutants are likely to arise in a *NOTCH1* mutant background, but whether the order of mutations impacts the selection of *TP53* mutants is unclear.

The 0.05 mm^2^ samples covered a total area 22.5 mm^2^ of epithelium from 7 donors. To determine if our findings were representative of a larger sample of tissue, we analysed 2 mm^2^ samples, comprising 48 cm^2^ of epithelium from 31 donors (**Supplementary Fig. 1b**, **Methods**). We first tested whether the mutations detected in the 0.05 mm^2^ and 2 mm^2^ sample sets were consistent with each other, finding strong correlation between the numbers of each type of mutation in the commonest mutated genes (r^2^ range 0.84-0.99, **Supplementary Tables 11-14**). The 2 mm^2^ samples showed that the proportion of mutant epithelium carrying nonsynonymous mutations in all positively selected mutant genes rose with age, so that the tissue approached saturation with mutant clones above the age of 70 (**Fig. 2a, Supplementary Table 14).** Mutant *NOTCH1* and *TP53* clones also expanded progressively, colonizing a median of 80% and 17% of epithelium in the over 70s respectively (**Fig. 2b, c, Supplementary Table 11**)^3, 7, 8^. Likewise, the proportion of nonsynonymous *NOTCH2* mutant epithelium rose with age, reaching a median of 5.7% in donors above age 70 (**Fig. 2d**). Comparison of mutant *NOTCH1*High and *NOTCH1*Low samples again showed that selection of *NOTCH2* was increased and mutant *TP53* decreased in the *NOTCH1*High samples (**Fig. 2e**). Furthermore, the VAF of *NOTCH2* mutations was increased across all age groups in *NOTCH1*High compared with *NOTCH1*Low samples (**Fig. 2f**). Conversely the VAF of *TP53* mutant clones was smaller in *NOTCH1*High samples in donors aged over 70 (**Fig. 2g**). Overall, these results are consistent with those from the 0.05 mm^2^ samples.

**Figure 2:**
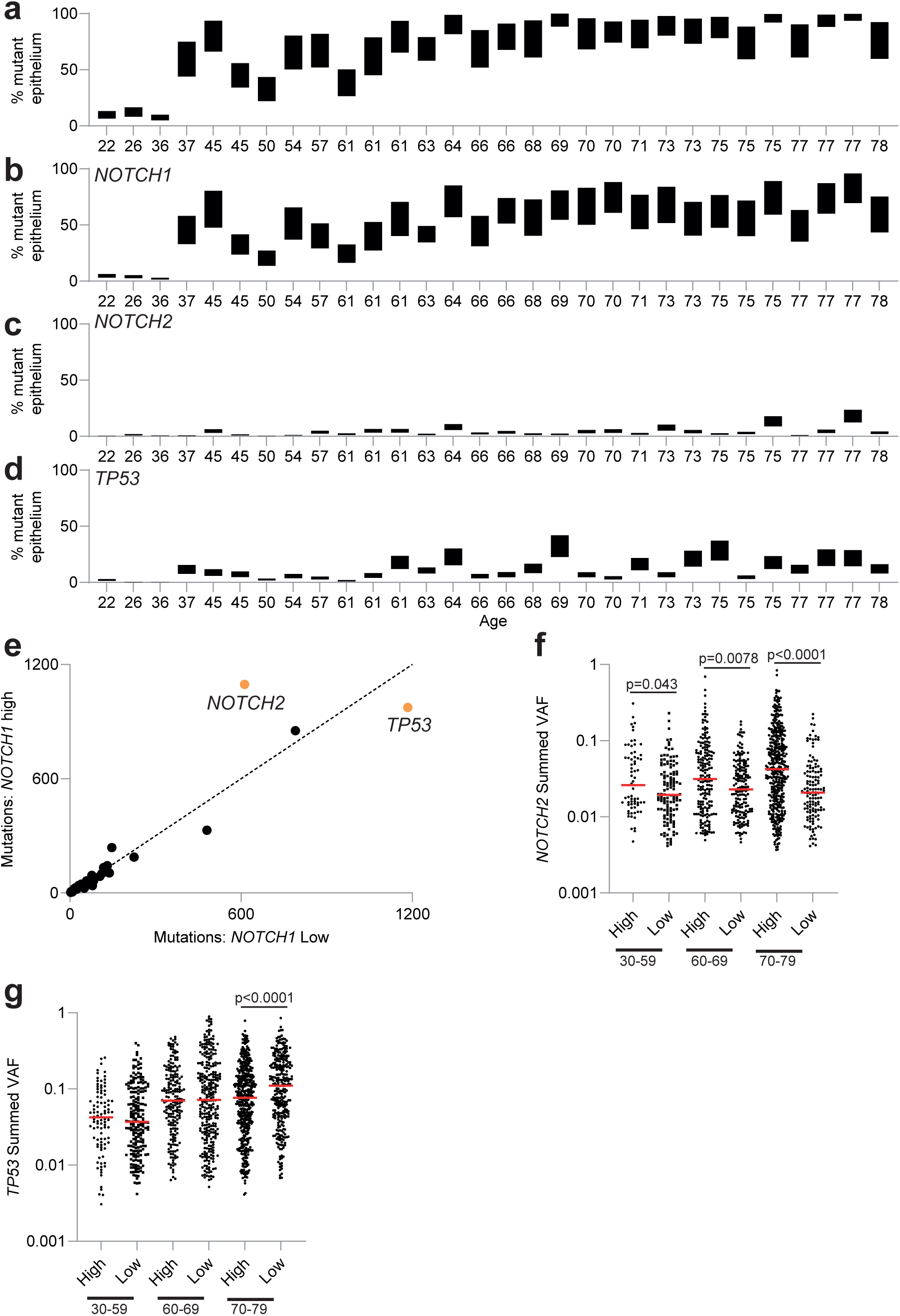
Mutational selection in 2 mm^2^ samples of human esophagus. **a-d** Proportion of mutant epithelium carrying nonsynonymous positively selected mutations. **a,** all positively selected mutants, **b**, *NOTCH1,* **c,** *NOTCH2,* and **d,** *TP53*. Upper and lower bounds reflect uncertainty in copy number and number of mutations per cell. Source Data: Supplementary Table 12. **e** Mutant selection in samples with different levels of *NOTCH1* mutation. 2 mm^2^ samples were sorted by the summed VAF of all *NOTCH1* mutants they carried into mutant *NOTCH1*High (*NOTCH1* summed VAF >0.4) and *NOTCH1*Low (*NOTCH1* summed VAF <0.4). Number of protein-altering mutations are plotted for 24 genes which have >100 mutations in either group (circles), orange indicates a significant difference. p values mutant: *NOTCH2* <0.0001, *TP53* 0.02, 2 tailed Z test. n samples *NOTCH1*High 1094, *NOTCH1*Low 1290. Source data: Supplementary Table 12. **f, g summed** VAF for nonsynonymous mutant *NOTCH2* clones (n=1552) (**f**), and nonsynonymous mutant *TP53* clones (n=2031) (**g**) in mutant *NOTCH1*High and *NOTCH1*Low samples in different age groups, 2 tailed Mann Whitney test. Source data: Supplementary Table 15.

We conclude aging human esophagus approaches saturation with positively selected mutant clones above the age of 60. In particular *NOTCH1* mutant cells colonize most of the tissue by age 60^3, 7, 8^. The continued acquisition of mutations (**Supplementary Fig. 1c**) suggests that the large population of *NOTCH1* mutant cells will acquire subclonal mutations over succeeding decades, so that epistatic interaction between *NOTCH1* and other mutants is feasible. The density of mutations in aged human esophagus also raises the possibility of spatial competition between adjacent mutant clones. Indeed, the distribution of mutant clone sizes is consistent with clonal competition in a near saturated landscape. However, it is not possible to determine the extent to which epistasis and/or spatial competition impact selection in *NOTCH1* mutant cells. We therefore turned to transgenic mouse models to explore the interactions of *Notch1*, *Notch2* and *Trp53* mutants in the esophagus.

### Fitness epistasis between mutant *Notch1* and mutant *Notch2* in mouse esophagus

We first investigated interactions between mutant *Notch1* and *Notch2*. Murine esophagus differs from humans in having a single, basal, proliferative cell layer, and fewer suprabasal cell layers, which undergo keratinized differentiation^27, 28^. However, as in human esophagus, there is no barrier to restrict the expansion of mutant clones within the basal cell layer^8, 29^. As a result, mutagenesis of mouse esophagus results in a dense patchwork of mutant clones resembling that of aged humans, with similar mutant genes under selection^4, 9^. As in humans, the most competitive mutant gene in murine esophagus is *Notch1*^7^. Losing the second allele of *Notch1* in a heterozygous mutant clone further increases competitive fitness, consistent with an excess of biallelic *NOTCH1* disruption in humans^7^. The mouse is thus a suitable model of mutant clonal fitness in the esophagus.

Based on our results in humans we hypothesised that *Notch2* mutants may have increased fitness in a *Notch1* mutant compared to a wildtype background. To test this, we aged mice in which both alleles of *Notch1* had been deleted in the esophagus (**Methods**). *Notch1^−/−^* epithelium was generated in *YFPCreNotch1^flox/flox^* mice, a transgenic line carrying a drug inducible *Cre* recombinase allele, and two conditional floxed *Notch1* alleles, as well as a neutral EYFP reporter in the *Rosa26* locus (**Fig. 3a, Methods**)^7^. *Cre* recombinase was activated by drug treatment of adult animals, resulting in deletion of the first exon of both alleles of *Notch1* (**Fig. 3b**). After ageing, a gridded array of 2 mm^2^ of esophageal epithelium samples was sequenced for 73 *Notch* pathway and cancer related genes (**Supplementary Table 15**)^7^. Uninduced littermates with a *Notch1* wild type esophagus were used as controls.

**Figure 3.**
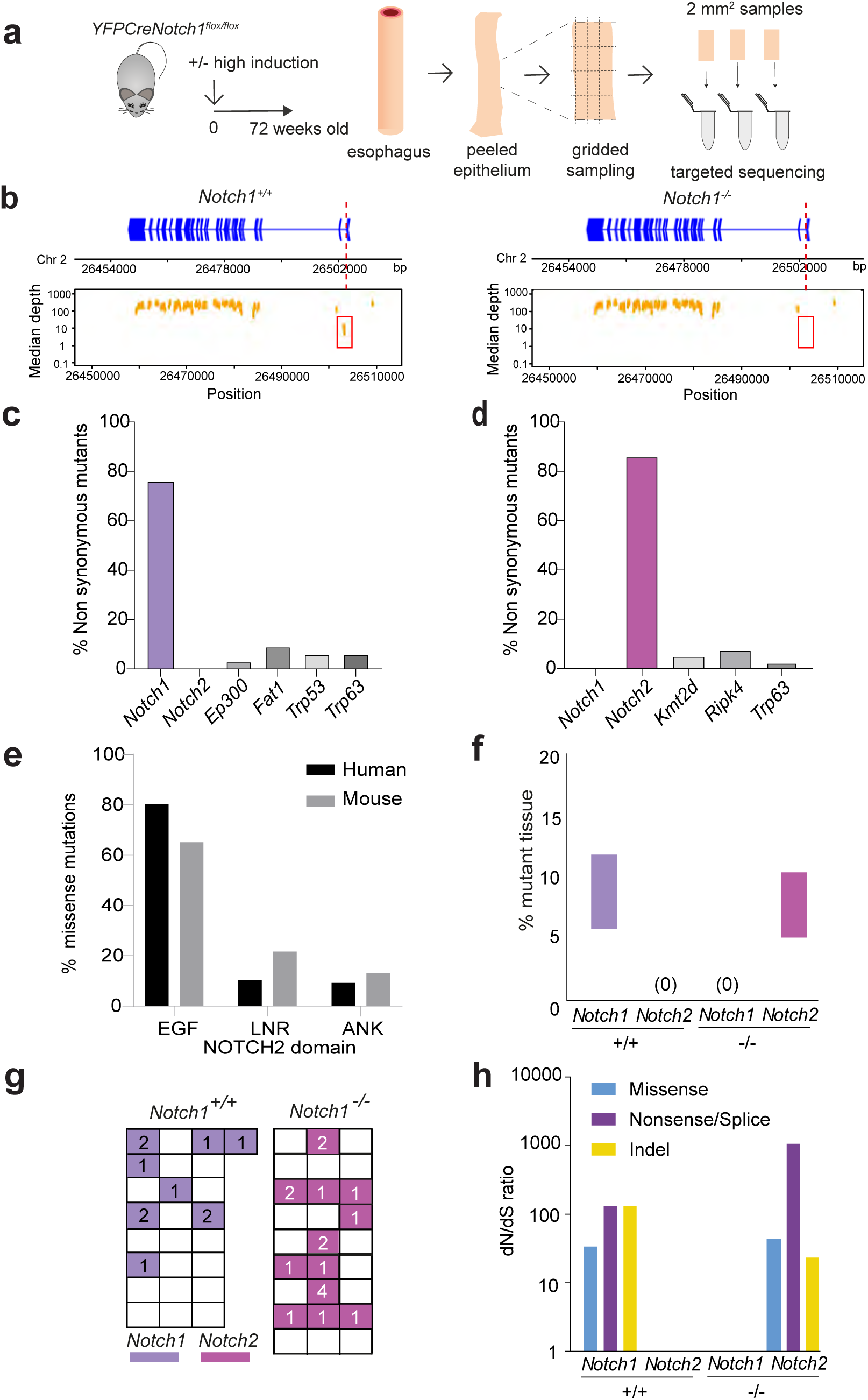
Selection of *Notch2* mutations in aging *Notch1^−/−^* mouse esophagus. **a** Protocol. Highly induced or uninduced control *YFPCreNotch1^flox/flox^* mice were aged. The esophagus was peeled and gridded into 2 mm^2^ samples that were processed for targeted sequencing (n= 50 or 51 total biopsies from 2 mice for each genotype, aged 72 weeks old ± 4 weeks). **b** Median depth of sequencing along *Notch1* gene in *Notch1^+/+^* and *Notch1^−/−^*esophagus from aging mice. Dashed red line and red box indicate site of *Cre* induced excision of the first exon of *Notch1*. **c, d Percentage** of all nonsynonymous mutations in mutant genes identified in aged *Notch1^+/+^* (**c**) and *Notch1^−/−^* esophagus (**d**) (total n = 33 and 42 respectively). 0 *Notch2* mutations were identified in *Notch1^+/+^* and 0 *Notch1* mutations in *Notch1^−/−^*esophagus. Source Data Supplementary Tables 19, 21. **e** Distribution of missense mutations affecting the EGF repeat (EGF), LNR and ANK domains of NOTCH2 mutations in humans (n=649) and mice (n= 23 from all samples). **f** Proportion by area of epithelium mutant for *Notch1* and *Notch2* in in control *Notch1^+/+^*(+/+) and induced *Notch1^−/−^*(−/−) esophagus, numbers in brackets indicate the number of nonsynonymous mutations of *Notch1* and *Notch2.* Source Data Supplementary Tables 19, 21. **g** Spatial distribution of mutations in a representative *Notch1* wild type and null esophagus, each box is a 2 mm*^2^* sample, numbers indicate number of specified mutations per sample, color indicates mutated gene. **h** dN/dS raMos for mutant *Notch1* and *Notch2* in control (+/+) and induced *Notch1^−/−^* (−/−) esophagus. Source data: Supplementary Tables 20, 22.

Mutant clones were identified in the aged epithelium both genotypes (**Fig. 3c, d, Supplementary Tables 16, 18**). The commonest nonsynonymous mutations in wild type epithelium were in *Notch1*. *Fat1*, *Trp53*, *Trp63* and *Ep300* mutants were also found, but no *Notch2* mutant clones were detected (**Fig. 3c**). However, in *Notch1^−/−^* epithelium, the most frequent nonsynonymous mutations were in *Notch2,* with rarer *Kmt2d*, *Ripk4* and *Trp63* mutants, and no *Notch1* mutants (**Fig. 3d**). Missense mutations in *Notch2* had a similar distribution across the protein to that seen in human esophagus (**Fig. 3e**). Similar areas of epithelium were colonised by *Notch1* and *Notch2* mutants in wild type and *Notch1* null esophagus respectively (**Fig. 3f**). Both *Notch1* and *Notch2* mutant clones were similarly distributed across the esophagus (**Fig. 3g**). A dN/dS analysis showed there was strong positive selection of *Notch1* mutants in wild type epithelium and of *Notch2* mutants in *Notch1* null esophagus. Collectively these results argue *Notch2* mutants are neutral in wild type tissue but become positively selected in the absence of *Notch1* (**Fig. 3h, Supplementary Tables 17,19**).

Mutant clones may be visualized by detecting areas of altered protein expression^7^. To determine if this approach was feasible for *Notch2* mutants, we examined *Notch2* expression in mouse esophagus by RNA sequencing, immune capillary electrophoresis and immunostaining of epithelial cyrosections (**Supplementary Fig. 5a-d, Supplementary Tables 20,21)**. *Notch2* was similar in both wild type and *Notch1^−/−^* epithelium. NOTCH2 was expressed at a low level in basal cells and higher levels in suprabasal cells whereas NOTCH1 was predominantly expressed in basal cells. These observations suggest immunostaining for NOTCH2 to detect *Notch2* mutant clones is feasible.

To test this approach, we aged a second cohort of mice with wild type and *Notch1^−/−^* esophagus and performed 3D confocal imaging of epithelial wholemounts immunostained for NOTCH2(**Fig. 4a**). Wholemounts were prepared by incubating esophageal epithelium with EDTA at 37 °C, which activated gamma secretase, triggering cleavage of the NOTCH2 transmembrane domain, allowing the cytoplasmic domain to migrate to the nucleus. (**Supplementary Fig. 5e**)^30^. In *Notch1* wild type epithelium, NOTCH2 staining was homogenous and nuclear, consistent with the absence of *Notch2* mutant clones detected by sequencing (**Fig. 4b**, **c**). However, in *Notch1^−/−^* tissue, there were scattered areas with decreased NOTCH2 staining (∼1/ mm^2^) (**Fig. 4b-d, Supplementary Table 22**). More rarely (∼ 0.1/ mm^2^), smaller areas with absent or altered protein localisation were observed located either within or adjacent to areas with decreased NOTCH2 staining (**Fig. 4c-e**; **Supplementary Fig. 5f; Supplementary Table 22**).

**Figure 4:**
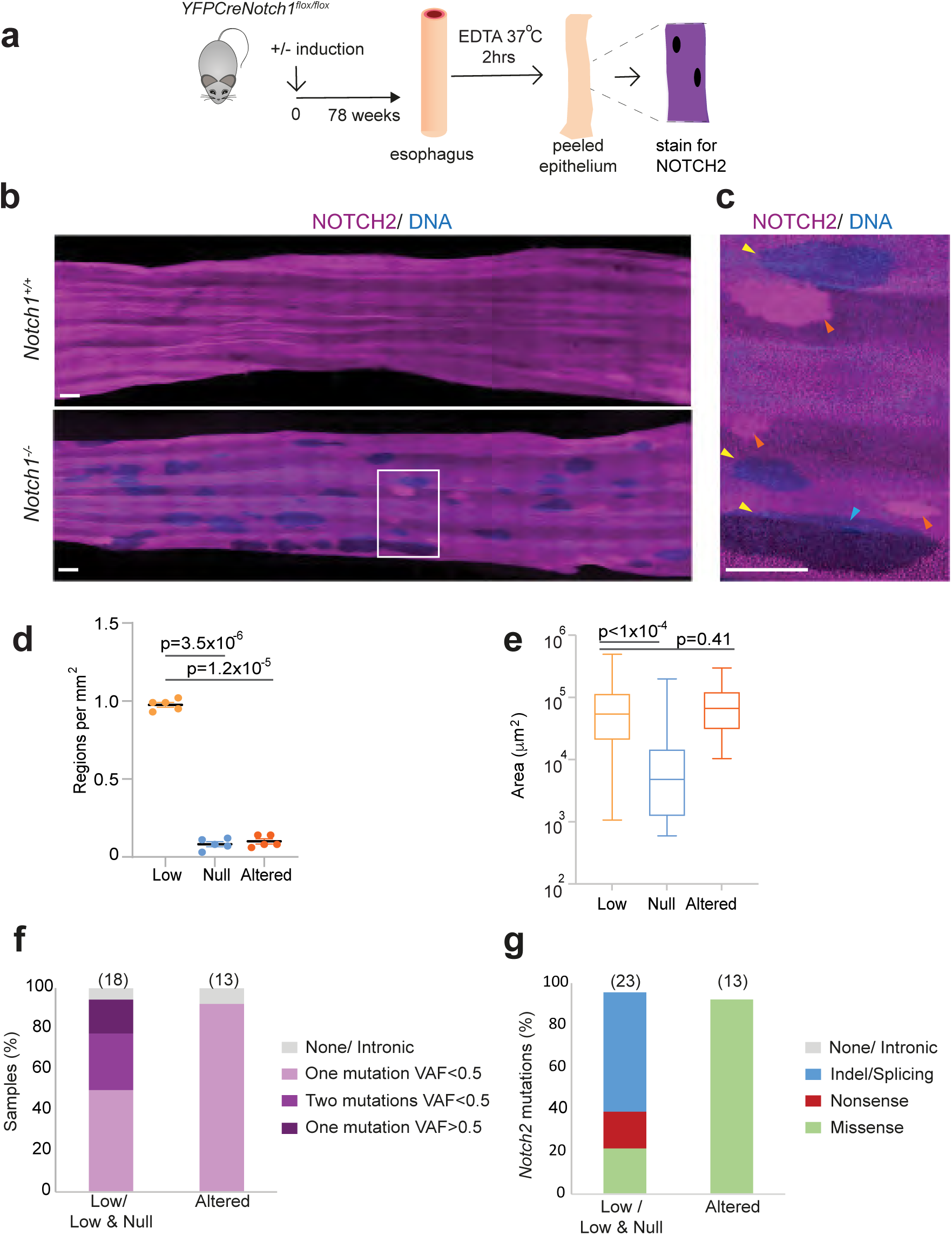
Localisation of NOTCH2 mutant clones by immunostaining and sequencing. **a** Protocol: Highly induced or uninduced control *YFPCreNotch1^flox/flox^* mice were aged. Whole mounts of the esophagus were then prepared by incubation in EDTA at 37 °C and stained for NOTCH2 and DNA. **b** Rendered confocal z stack images showing top-down views of representative immunofluorescence staining for NOTCH2 (magenta) and DNA (blue) in wholemount epithelium from aged wild type and *Notch1^−/−^* mice. Images are representative of entire esophagus from 5 *Notch1^+/+^* and 11 *Notch1^−/−^* animals. Scale bars, 500µm. **c** Magnified image from area indicated by white box in **b**. Yellow arrowheads, low intensity staining; orange arrowheads, altered positive staining; blue arrowheads, null staining areas. Scale bar, 500µm**. d** Number of regions with altered NOTCH2 staining per mm^2^, each dot indicates average per mouse, n=5 mice, p values from repeated measures ANOVA with post hoc Bonferroni correction. Source Data: Supplementary Table 25. **e** Area of regions with altered NOTCH2 staining. n=277 low intensity, 24 null, and 29 altered positive foci from 5 mice. Kruskal Wallis test, post hoc Dunn’s comparison. Source Data: Supplementary Table 25. **f** Proportion of samples from areas of altered staining carrying one mutation with VAF<0.5, two mutations with VAF<0.5, one mutation with VAF>0.5 or with intronic or undetected *Notch2* mutations. Number of samples is shown in brackets. Low and null refers to samples containing both low and null staining areas **g.** Proportion of *Notch2* missense, nonsense, indel/splicing or intronic/undetected mutations in samples with aberrant NOTCH2 staining. Number of mutations is shown in brackets. Low and null refers to samples containing both low and null staining areas Source data: Supplementary Tables 26,27.

Next, we performed targeted sequencing of areas with altered NOTCH2 staining, collected with a 0.05 mm^2^ biopsy punch (**Supplementary Fig. 5g**). This revealed the presence of near clonal nonsynonymous mutations *Notch2* (average VAF 0.31) (**Fig. 4f**, **Supplementary Tables 23,24**). Most areas with reduced NOTCH2 staining harbored a single *Notch2* mutation consistent with a heterozygous mutant clone (**Fig. 4f**). Samples containing null staining areas contained either a single mutation with VAF>0.5 or two mutations consistent with biallelic *Notch2* mutation/LOH (**Fig. 4f**). Areas with altered NOTCH2 staining pattern carried missense mutants in domains that disrupt NOTCH2 protein function (**Fig. 4g**). We conclude that heterozygous *Notch2* mutation increases clonal fitness in a *Notch1* null background. Clones with biallelic *Notch2* disruption emerge from heterozygous *Notch2* mutant clones, arguing the complete loss of *Notch2* further increases mutant fitness.

To investigate how *Notch2* mutation alters cell behavior we performed lineage tracing with 5-ethynyl-2’-deoxyuridine (EdU). EdU labels cells in S phase and can be used to track the fate of proliferating cells and their progeny by using 3D imaging (**Fig. 5a**). Aged mice with a *Notch1* null esophagus were given a dose of EdU either one or 48 h prior to sacrifice. We first compared areas of normal or altered NOTCH2 staining in the esophagus of animals culled 1 hr post EdU administration (**Fig. 5b**-**d**). We found no significant differences in basal cell density or the proportion of EdU positive basal cells between the different staining areas (**Fig. 5e, f, Supplementary Tables 25,26**). Next, we analysed animals imaged 48 h post EdU labelling, when some of the EdU labelled cells have differentiated and migrated into the first suprabasal cell layer (**Fig. 5g-i**). The proportion of these EdU+ suprabasal cells reflects the rates of differentiation and migration. This was significantly reduced in areas with absent compared with reduced or normal NOTCH2 staining (**Fig. 5j, Supplementary Table 27**). We concluded that of loss of NOTCH2 on a NOTCH1 null background reduces the rate of differentiation. This phenotype is also seen when NOTCH signalling is blocked by overexpression of dominant negative mutant *Maml1*^29^. A spatial Moran type simulation suggested decreased cell loss through slower differentiation is a potential mechanism for cells with biallelic *Notch2* loss to gain a fitness advantage over their heterozygous *Notch2* mutant neighbors (**Fig. 5k)**.

**Figure 5:**
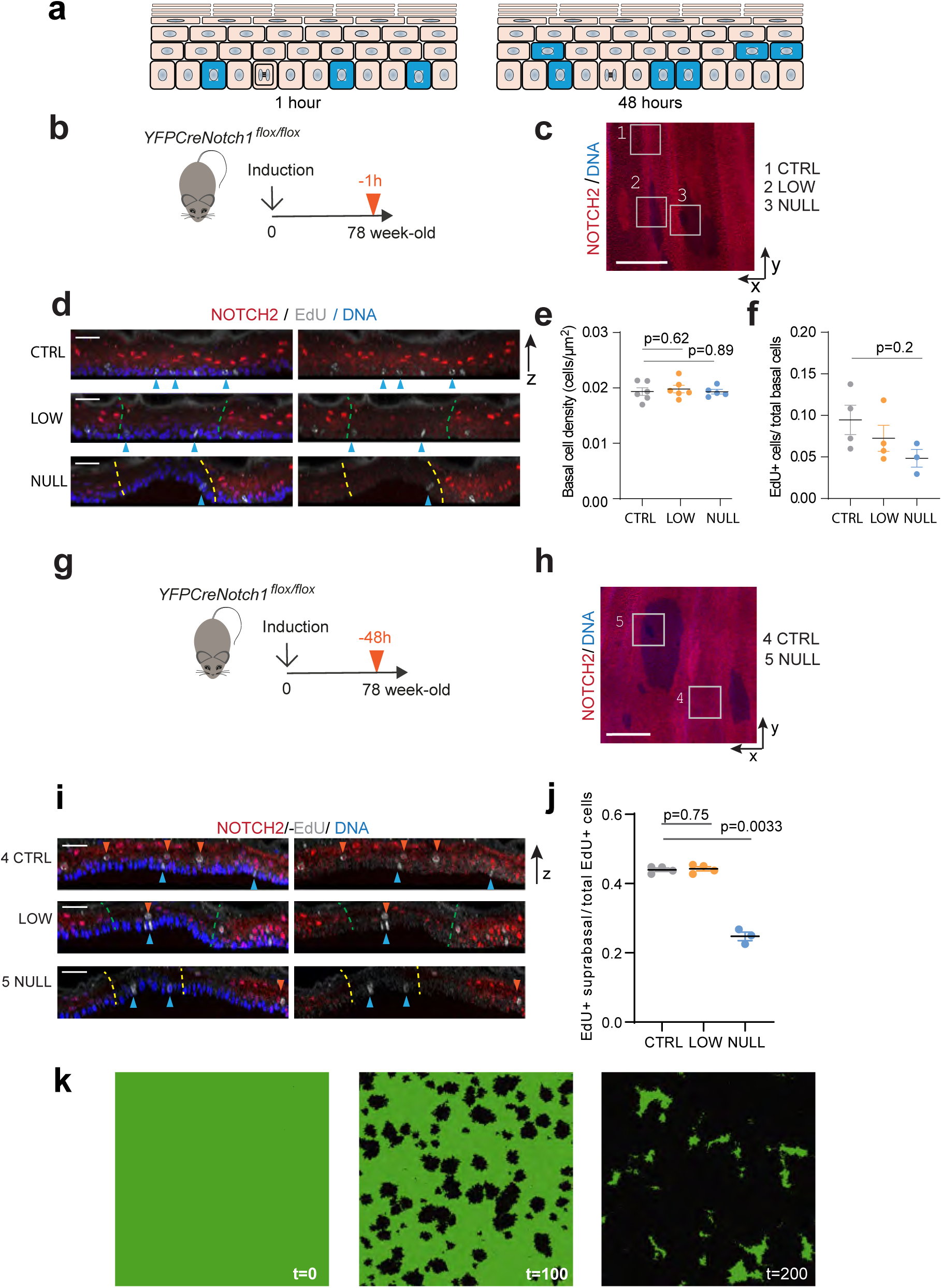
Cell dynamics in *Notch2* mutant areas. **a.** EdU labelling experiments in mouse esophageal epithelium. EdU is injected 1 or 48 hours before culling. At 1h, EdU labels S phase cells in the proliferating basal cell layer (Blue). At 48 h, cells have divided and labelled cell pairs are seen in the basal and first suprabasal cell layers. The ratio of EdU+ suprabasal/total EdU+ cells reflects the rate of stratification. **b** Protocol. Highly induced *YFPCreNotch1^flox/flox^* mice were aged for up to 78 weeks of age and injected with EdU 1 h before esophageal tissue collection. **c, d** Representative rendered projected confocal 3D z stack images of epithelial wholemounts stained with NOTCH2 (red), DNA (blue) and EdU (grey). Focal areas of NOTCH2 staining were analysed, 1 normal (CTRL), 2 reduced (LOW), 3 absent (NULL) **c**, top down (x, y) and **d** lateral (z) views of epithelium. In **d**, blue arrows indicate EdU+ basal cells, Dashed lines delineate NOTCH2 low intensity (green) or null (yellow) staining regions. Scale bars: **c**, 500 µm, **d**, 30 µm. **e,** Basal cell density in NOTCH2 normal (CTRL), low, and null staining areas. Mean ± s.e.m., each dot represents a mouse. CTRL, n = 6 mice (total 7477 cells); LOW, n = 6 mice (6118 cells); NULL, n=5 mice (2454 cells). Repeated measures Anova, p-value against control condition from post hoc Bonferroni correction. Source data: Supplementary Table 25. **f,** Proportion of EdU+ basal cells in mice injected with EdU 1 hour previously. Mean ± s.e.m., each dot represents a mouse, CTRL, n = 4 mice (6295 cells); LOW, n = 4 mice (4973 cells); NULL, n= 3 mice (1294 cells). 1 way ANOVA. Source data: Supplementary Table 26. **g** Protocol. Highly induced *YFPCreNotch1^flox/flox^* mice were aged for up to 78 weeks of age and injected with EdU 48 h before esophageal tissue collection. **h, i** Representative rendered projected confocal 3D z stack images of epithelial wholemounts stained with NOTCH2 (red), DNA (blue) and EdU (grey). Focal areas of NOTCH2 staining were analysed, normal (CTRL, 4), reduced (LOW), absent (NULL, 5) in top down (x, y), **h**, and lateral, (z), **i**, views of epithelium. Blue arrows indicate EdU+ basal cells, orange arrows EdU+ suprabasal cells. Dashed lines delineate NOTCH2 low intensity (green) or null (yellow) staining regions. Scale bars, **h**, 500 µm, **i**, 30 µm. **j** Ratio of EdU+ suprabasal: total EdU+ cells. Mean ± s.e.m., each dot represents a mouse, CTRL, n =4 mice (2316 total cells); LOW, n = 4 mice (1989 cells); NULL, n= 3 mice (451 cells). Repeated measures ANOVA with Bonferroni post hoc correction. Source data: Supplementary Table 27. **k** Stills from a spatial Moran like simulation embodying the observations above in of *Notch2^−/−^* (black) in a heterozygous *Notch2* mutant background (green), time units are cellular generations. See **Video 1** and **Supplementary Note** for details.

In summary, these results argue *Notch2* mutants compete neutrally in a wildtype background and hence do not generate detectable mutant clones. However, in a *Notch1* null esophagus, epistasis between *Notch1* and *Notch2* increases the fitness of the mutant clones leading to clonal expansion when a spontaneous *Notch2* mutation occurs. The emergence of areas lacking NOTCH2 protein with a decreased rate of differentiation suggests that biallelic *Notch2* loss further increases fitness.

### Selection of mutant *Trp53* and *Notch1* in murine esophagus

Next, we studied interactions between *Trp53* and *Notch1* mutant clones in mouse esophagus. Individually both *Notch1* and *Trp53* mutation drive clonal expansions in wild type esophagus^5,7^. To address the relative fitness and interactions of single and double *Notch1* and *Trp53* mutants within the same tissue, we used a second mouse model, Cyp*1a1Cre^ERT^Trp53^R245W-GFP/Wt^ Notch1^flox/flox^* mice. These animals carry conditional alleles of both *Notch1* and *Trp53*. Treatment with low doses of inducing drugs results in transient expression of *Cre* recombinase^27^. Levels of induction were titrated to give expression of one or both mutant alleles in scattered individual progenitor cells. Clones were detected by immunostaining for GFP, which reports mutant *Trp53 expression,* and NOTCH1 to detect *Notch1* deletion in epithelial wholemounts (**Fig. 6a)**^5, 7^.

**Figure 6:**
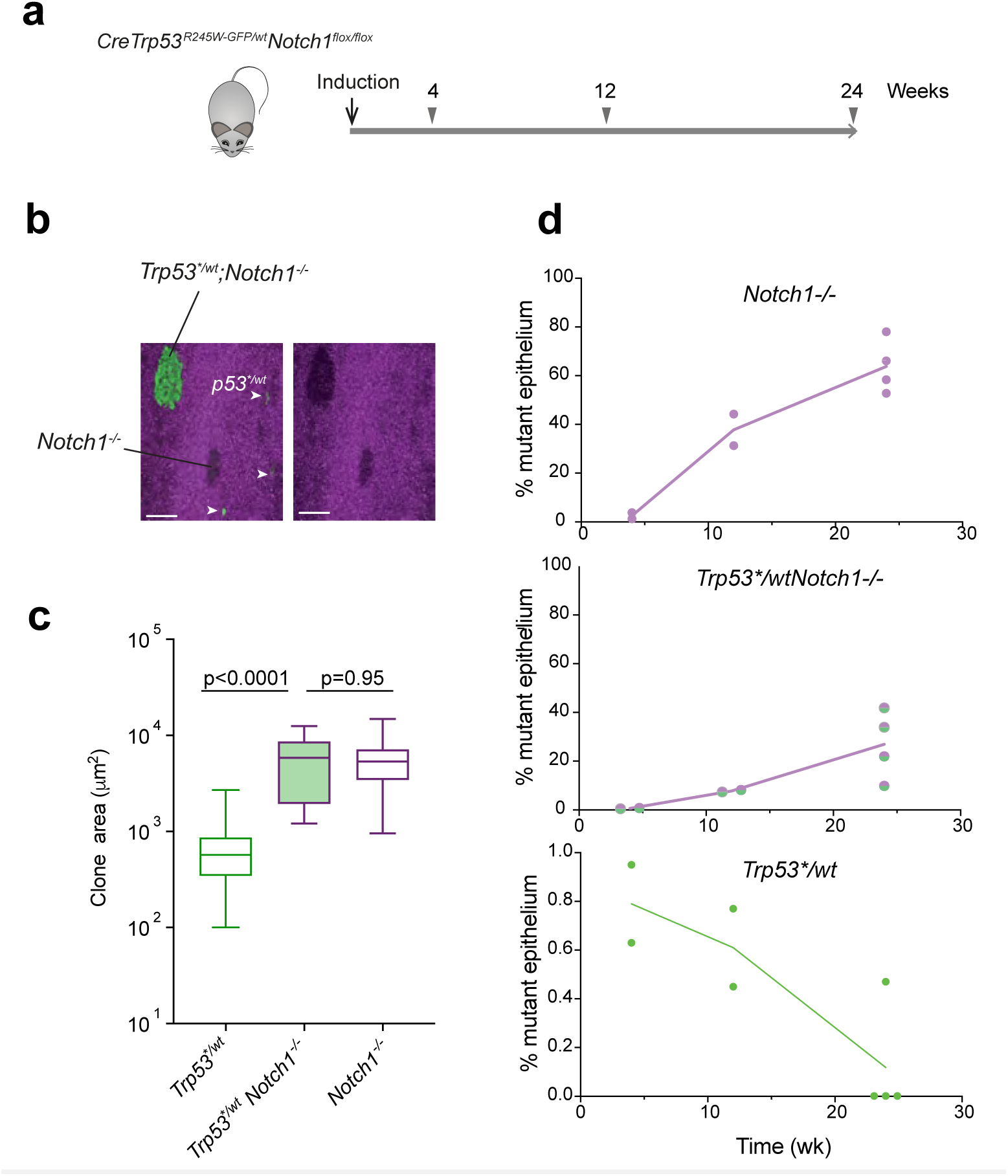
*Notch1* and *Trp53* mutants in mouse esophagus. **a**: Protocol: *Cyp1a1Cre^ERT^Trp53^R245W-GFP/Wt^ Notch1^flox/flox^ mice* were induced, and esophageal samples were collected at 4, 12 and 24 wk later (triangles). n = 2 mice at 4 and 12 wk, n = 4 mice at 24 weeks. **b**: Representative rendered confocal z stack image showing top-down view of the basal layer of an epithelial wholemounts from the esophagus 4 wk amer clonal induction. Green is GFP linked to mutant *Trp53^R245W^*(*p53^*/wt^*) expression, clones indicated by white arrows. *Notch1^−/−^* clones detected by lack of immunostaining for NOTCH1, purple. *p53^*/wt^; Notch1^−/−^* clones express GFP but not NOTCH1. Scale bar, 100 μm. **c**: Clone size distribution at 4 wk Mme point. Area of each clone was measured and plo(ed. Box indicates quartiles, line median and whiskers range of clone size. *p53^*/wt^*, 330 clones, *p53^*/wt^: Notch1^−/−^* 29 clones, and *Notch1^−/−^*155 clones, from two mice. *p* value, 2 tailed Mann-Whitney test. Source data: Supplementary Table 28. **d**: Proportion of epithelium with *p53*/wt*, Notch1*^−/−^*, or *p53^*/wt^; Notch1^−/−^* mutations by area. n mice = 2 at 4 wk, 2 at 12 wk and 4 at 24 wk. Source data: Supplementary Table 29.

We induced *Cre* and prepared esophageal wholemounts at 4-, 12– and 24-wk post induction (**Fig. 6a**). At 4 wk after induction, clones surrounded by wild type cells were observed. Single mutant *Trp53^R245/wt^* clones were substantially smaller than that of either single mutant *Notch1^−/−^* or double mutant *Notch1^−/−^Trp53^R245W/wt^* clones in the same esophagus (median fold difference in clone size 10 and 9.4 respectively) (**Fig. 6b, c, Supplementary table 28)**. There was no significant difference between the size of *Notch1^−/−^* and *Notch1^−/−^Trp53^R245W/wt^* clones (**Fig. 6c)**. We concluded that *Notch1* loss has a dominant effect on the competitive fitness of double mutant clones, the addition of a *Trp53* mutation conferring no detectable benefit.

By 12 wk post induction, clones had expanded and many had collided, so we instead measured the area of epithelium mutant for *Trp53*, *Notch1* or both (**Fig. 7d, Supplementary Table 29**). There was 7-fold decrease in the average area of *Trp53* mutant epithelium at 24 wk compared with 4 wk, while the area of both single mutant *Notch1^−/−^* and double mutant *Notch1^−/−^ Trp53^R245W/wt^* epithelium rose significantly (Kruksal-Wallis test, p=0.0049 and 0.014 respectively). In three of 4 animals at 24 wk there was no remaining *Trp53* mutant epithelium as the entire tissue was occupied by *Notch1^−/−^* and *Notch1^−/−^Trp53^R245W/wt^* mutant cells (**Fig. 6d**). These findings are consistent with spatial competition in which the less fit *Trp53^R245W/wt^* mutants have been displaced from the tissue by the expansion of fitter *Notch1^−/−^* and *Notch1^−/−^ Trp53^R245W/wt^* clones^4^.

**Figure 7:**
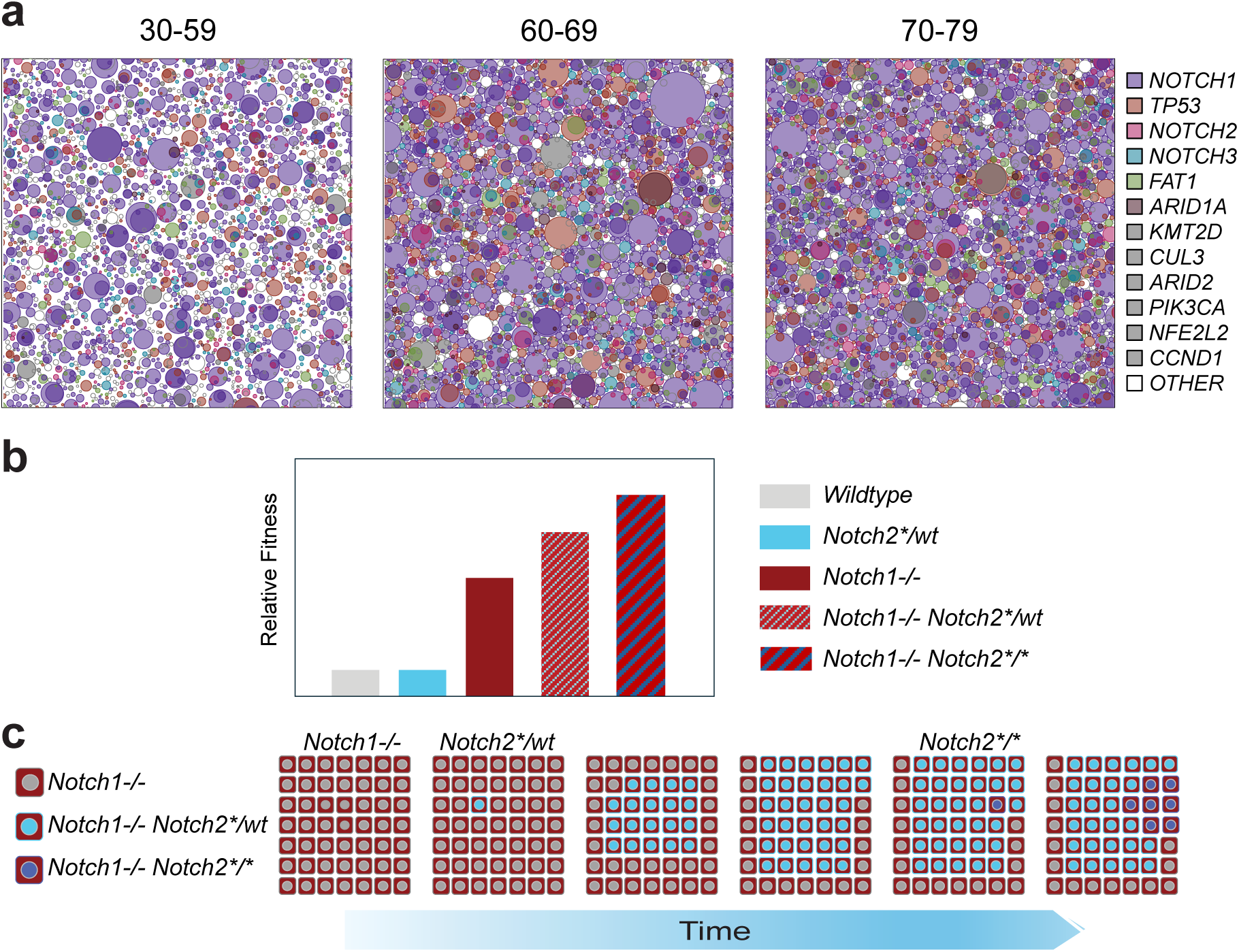
*Notch1 is* epistatic with *Notch2*. **a** Representative visualization of mutations in human 2mm^2^ samples. Top-down views of nonsynonymous mutant clones for mutant genes under positive selection in 9 cm^2^ of epithelium in the age groups shown. 2mm^2^ samples were randomly selected and mutations displayed as circles, randomly distributed in space. Sequencing data was used to infer the size and number of clones and, where possible, the nesting of sub-clones. **b** Relative fitness of wild type, *Notch1*^−/−^ 7 and *Notch2* mutants inferred from results in mouse esophagus. *Notch2* mutants appear neutral in a wildtype background, but subclonal *Notch2* mutation enhances fitness in *Notch1^−/−^* cells. Loss of the second *Notch2* allele further increases fitness in a *Notch1^−/−^* background. This results in clonal expansions of *Notch2* mutants as shown in **c**.

## Discussion

Our observations show that in the densely mutated aged human esophagus, the trajectory of subsequent mutational selection is altered by the extensive colonization of the tissue by *NOTCH1* mutants that occurs by age 60 (**Fig. 7a**). Mutant *NOTCH2* selection is increased and *TP53* mutations decreased in *NOTCH1* mutant tissue. *NOTCH2* mutations predominantly arise within *NOTCH1* mutant clones. Double mutant *TP53/NOTCH1* mutant clones are rarer, but their existence shows that the two mutations are not synthetically lethal. However, there are too few *TP53/NOTCH1* mutants for reliable inference of the order of mutations. Taken alone, sequencing normal human esophagus reveals altered selection but cannot resolve the underlying mechanism(s).

Mouse models give insight into the basis of altered selection. In ageing wild type esophagus, *Notch2* mutants compete neutrally with adjacent wild type cells and fail to generate clones detectable by sequencing or immunostaining. However, when we aged mice with a *Notch1^−/−^* esophagus, heterozygous *Notch2* mutant clones emerged, indicative of epistasis. The emergence of rarer areas with absent NOTCH2 staining containing two *Notch2* mutants/LOH argues loss of the second *Notch2* allele further increases fitness above that of heterozygous *Notch2.* This is at least in part due to a decreased rate of cell loss through stratification in areas with biallelic *Notch2* disruption. Collectively these observations are consistent with fitness epistasis between *Notch1* and *Notch2* mutants in *Notch1^−/−^* esophagus, with fitness rising as total *Notch* gene dosage decreases (**Fig. 8 b, c**). The mouse studies argue that epistasis explains the observation that almost all *NOTCH2* mutations in humans occur within clones carrying *NOTCH1* mutation(s) (**Fig. 1g**).

**Figure 8:**
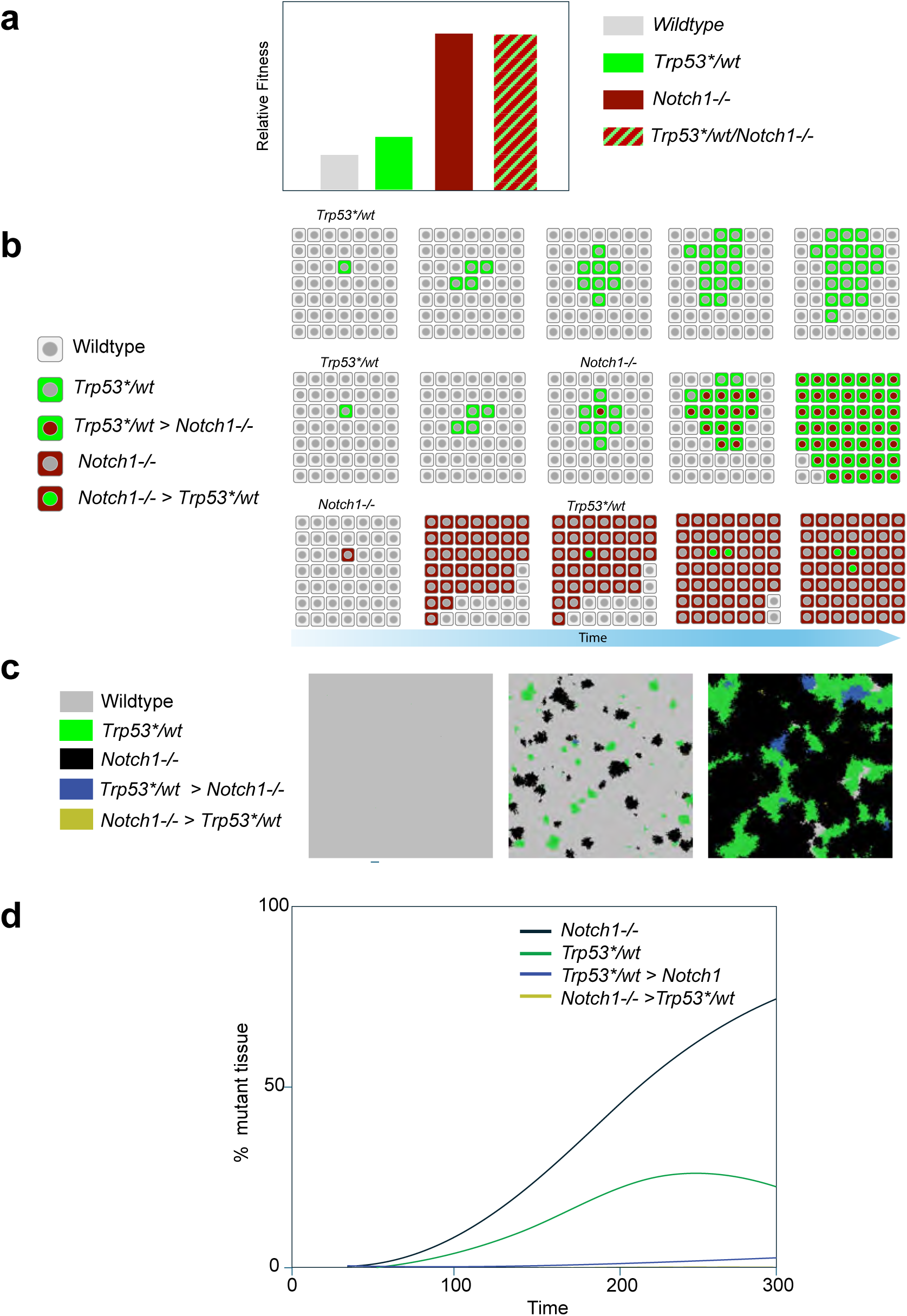
Fitness and selection of double mutant *Trp53* and *Notch1* clones. **a** Relative fitness of mutants, inferred the size of clones expanding in a wildtype background (**Fig. 6b**). **b** Hypothesis: order of mutations and fitness of neighboring cells impact clonal expansion. Mutant *Trp53*\* cells outcompete surrounding wild type cells. *Notch1* mutation (*Notch1\**) within a *Trp53*\* clone increases fitness, so a double mutant clone expands rapidly outcompeting both wild type and single mutant cells. In contrast, single mutant *Notch1* null clones expand more rapidly than *Trp53\** mutants due to their higher fitness. The occurrence of a *Trp53*\* mutation within a *Notch1* null clone has minimal effect on fitness, so the double mutant clone will compete (near) neutrally with its neighbors. **c, d** Simulation of the inferred fitness of *Trp53* mutant, *Notch1^−/−^* and double mutant clones. **c** Snapshots of simulated clonal expansion over time. A spatial-Moran process is implemented with a random mutation process altering 1 in 10,000 cells every generation, selecting *Trp53* mutation or *Notch1* loss 522 with equal probability. Following experimental observations, *Trp53* mutants are less fit than *Notch1^−/−^* clones in a wildtype background, and the double mutant has the same fitness *Notch1^−/−^* clones. Both *Trp53* and *Notch1^−/−^* mutant clones expand initially, with *Trp53* mutant clones regressing when they collide with fitter *Notch1^−/−^*clones. Double mutant clones where *Trp53* muta*on occurs first are more able to expand and persist due to their fitness increase compared with *Trp53* mutant neighbors. In contrast, where *Notch1* loss occurs first there is no increase in fitness compared with neighboring *Notch1* null cells. Wild type background is magenta, *Notch1^−/−^* clones are black, *Trp53* mutant green, and the double mutants yellow or blue depending on whether Notch1 or *Trp53* mutation occurs first. T represents tissue generations. **d** Colonisation of wild type esophagus by different mutants over time in model shown in **c** *Trp53* > *Notch1* and *Notch1* > *Trp53* indicate double mutants in which *Trp53* or *Notch1* occurred first respectively. See **video 2** and **Supplementary Note** for details.

Turning to the interaction of *Trp53* and *Notch1,* there was no detectable fitness epistasis between the two mutants in mouse esophagus, judging by the size of single and double mutant clones expanding in a wildtype background (**Fig. 8a**). However, our results suggest that the order of mutations and the genotype of surrounding cells may have a substantial effect on the fate of the double mutant clones (**Fig. 8b**). If a cell within a *Trp53* mutant clone acquires a *Notch1* mutation, the fitness of double mutant cells is increased substantially compared to the surrounding *Trp53* mutants. In consequence the double mutant clone will be likely to outcompete the surrounding *Trp53* single mutant and wild type cells and expand substantially. In contrast, a *Trp53* mutant within a *Notch1* null area gains competes at or near neutrally with the surrounding *Notch1* null cells and so is less likely to expand and persist (**Fig. 8b**).

As well as such ‘neighbor fitness’ effects, spatial competition also impacts the survival and expansion of mutant clones (**Fig. 8c, d**). When expanding clones collide, *Trp53* single mutants are displaced from the tissue by either *Notch1* null or *Trp53, Notch1* double mutant clones. How might these processes interact to influence the proportions of mutant cells? We explored this in a simple spatial-Moran process model in which a random mutation occurring at a constant rate every cellular generation, generates a *Trp53* or *Notch1* mutation with equal probability (**Fig. 8c**). The model encompassed our experimental observations. *Trp53* mutations are less fit than *Notch1* mutant clones in a wildtype background, but double mutants have the same fitness as clones mutant for *Notch1* alone. In the model, as in mice, *Trp53* and *Notch1^−/−^* single mutant clones expand initially but then regress when they collide with fitter single or double *Notch1* mutant clones. Double mutant clones in which *Trp53* mutation occurs first expand but those in which *Trp53* occurs second compete neutrally. As the simulation continues, the proportions of *Notch1* and *Trp53* single mutant and double mutant epithelium evolve to generate a landscape resembling that of humans (**Fig. 8d**).

What are the potential implications of these findings for carcinogenesis? The colonization of the esophagus by *NOTCH1* increases the proportion of *NOTCH2* mutations in aged esophagus. However, the proportion of *NOTCH2* mutant epithelium is similar to that in ESCC, arguing *NOTCH2* mutation does not majorly increase the likelihood of malignant transformation (**Supplementary Fig 1a, Fig 2c**). Our findings argue the spread of *NOTCH1* mutant clones may limit the expansion of the *TP53* mutant *NOTCH1* wild type clones from which most ESCC originate. What of the risk of transformation of *TP53, NOTCH1* double mutant epithelium? An upper bound estimate of the proportion of double mutant cells in over 60s is 7% (33/451 samples, **Fig. 2b**) compared with 12% (11/89) of ESCC (**Supplementary Fig. 1a**, p=0.13, Fisher’s exact test). This suggests that the oncogenicity of *TP53, NOTCH1* double mutant epithelium may be attenuated compared with *TP53* mutant *NOTCH1* wild type tissue. These observations in squamous esophageal carcinogenesis have an interesting parallel in the somatic evolution of metaplastic Barrett’s esophagus (BE) in the distal esophagus and gastroesophageal junction. The fitness of *TP53* mutant clones in this glandular epithelium is reduced in cells which have lost CDKN2A, lowering the population of *TP53* mutants^31^.

It is well established that ‘order matters’ for the selection of mutants in cancer genomes^32, 33, 34, 35, 36^. We conclude that this principle also operates in the normal esophagus. Epistasis, the fitness of adjacent cells and competition between colliding clones all influence the trajectory of somatic evolution. In humans the colonization of most of the esophagus by highly fit *NOTCH1* mutants decreases the selection of less fit oncogenic *TP53* mutants. More broadly, dominant mutations that colonize extensive areas in normal tissues may influence selection of oncogenic mutants in epithelial carcinogenesis and the risk of cancer development.

## Acknowledgements

We thank Esther Choolun and Tom Metcalf for assistance with *in vivo* experiments, and Wellcome Sanger Institute RSF facilities for technical support.

This work was supported by grants from the Wellcome Trust to the Wellcome Sanger Institute (098051, 296194, and 108413/A/15/D) and Cancer Research UK Programme Grants to P.H.J. (C609/A17257 and C609/A27326). BAH is supported by a Royal Society URF grant (UF130039).

## Author Contributions

J.C.F., E.A, K.M., and C.K. performed the experiments. C.K., R.S., S.B., S.H.O., F. W-L., and J.C.F. processed and analyzed sequencing data. I.A. and M.W.J.H performed statistical analysis supervised by B.H. B.H generated simulations. P.H.J. supervised the project and wrote the manuscript with input from C.K., E.A. R.S., B.H., and J.C.F.

## Supplementary Material

The Supplementary Note follows methods.

## Supplementary Figures and legends

**Supplementary Figure 1:**
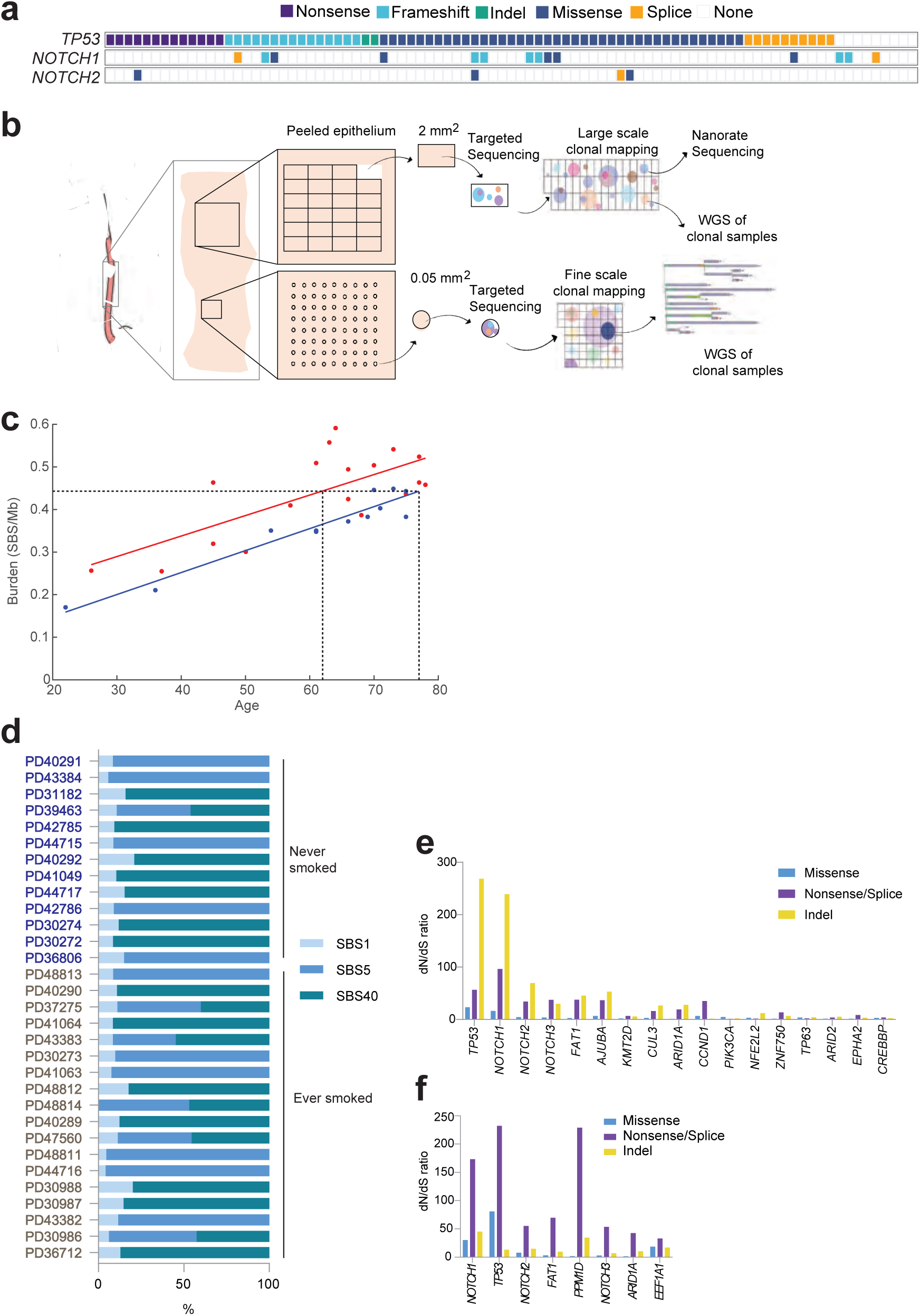
Mutational burden, mutational signatures and mutant selection in aging esophagus. **a** Single nucleotide variants in ESCC from^10^ **b** Sequencing Protocol. Normal epithelium of the middle third of the esophagus from transplant donors was sampled by taking arrayed samples of 2 mm^2^ and 0.05 mm^2^ for targeted sequencing. A subset of clonal samples was analysed by whole genome sequencing (WGS). Mutational burden was determined from NanoSeq analysis of a 2 mm^2^ sample from each donor**. c** Mutational burden (single base substitutions/megabase) in ever smokers (red), and never smokers (blue) determined by Nanoseq. Lines show linear correlation of burden with age (smokers R=0.714, p=0.00128; non-smokers R=0.95 p=0.00001). Dashed lines projected to the x-axis show age difference between smokers and non-smokers to reach the same mutational burden projected to the y-axis. Source data: Supplementary Table 3. **d** Mutational signatures of single base substitutions from Nanoseq. Source data: Supplementary Table 3. **e**, **f** Positive selection of mutant genes. dN/dS ratios of missense (Turquoise), nonsense/splice (purple) and indels (yellow) for positively selected genes in 2 mm^2^ samples, **e**, and 0.05 mm^2^ samples, **f**. Neutral, unselected genes have dN/dS=1. dN/dS ratios calculated with dNdScv. (q<0.01). Source data: Supplementary Table 4.

**Supplementary Figure 2:**
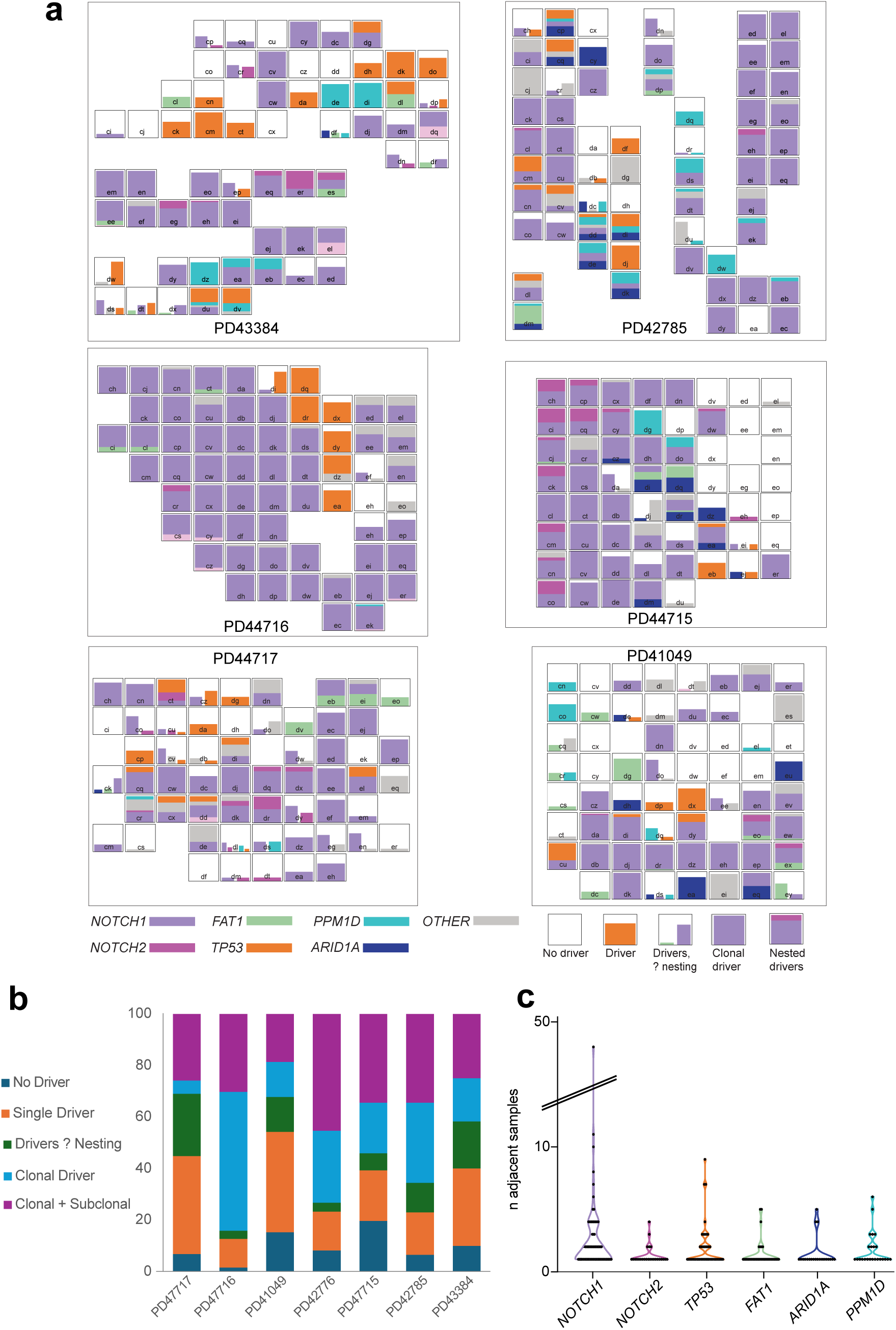
Mapping mutations in 0.05 mm^2^ samples. **a** Targeted sequencing of tissue shown in **a** for 324 cancer associated genes. Bar charts show proportion of sample mutant for positively selected mutants shown. Position of bar chart corresponds to spatial location of sample; letters indicate sample identity. Bar height indicates proportion of mutant tissue derived from variant allele frequency. Stacked bars indicate nested mutations, side-by side bars indicate nesting is unknown, color indicates mutant gene, ‘other’ indicates mutation in another positively selected mutant gene. Source data: Supplementary Table 7. **B** Percentages of samples carrying driver(s) in donors shown in **Fig.1b** and panel **a**. Number of samples 451. Donor ages ranged from 61-75 (Supplementary Table 1). **c** Number of adjacent samples spanned by clonal mutations of the positively selected mutant genes shown across all donors Source data: Supplementary Table 7.

**Supplementary Figure 3:**
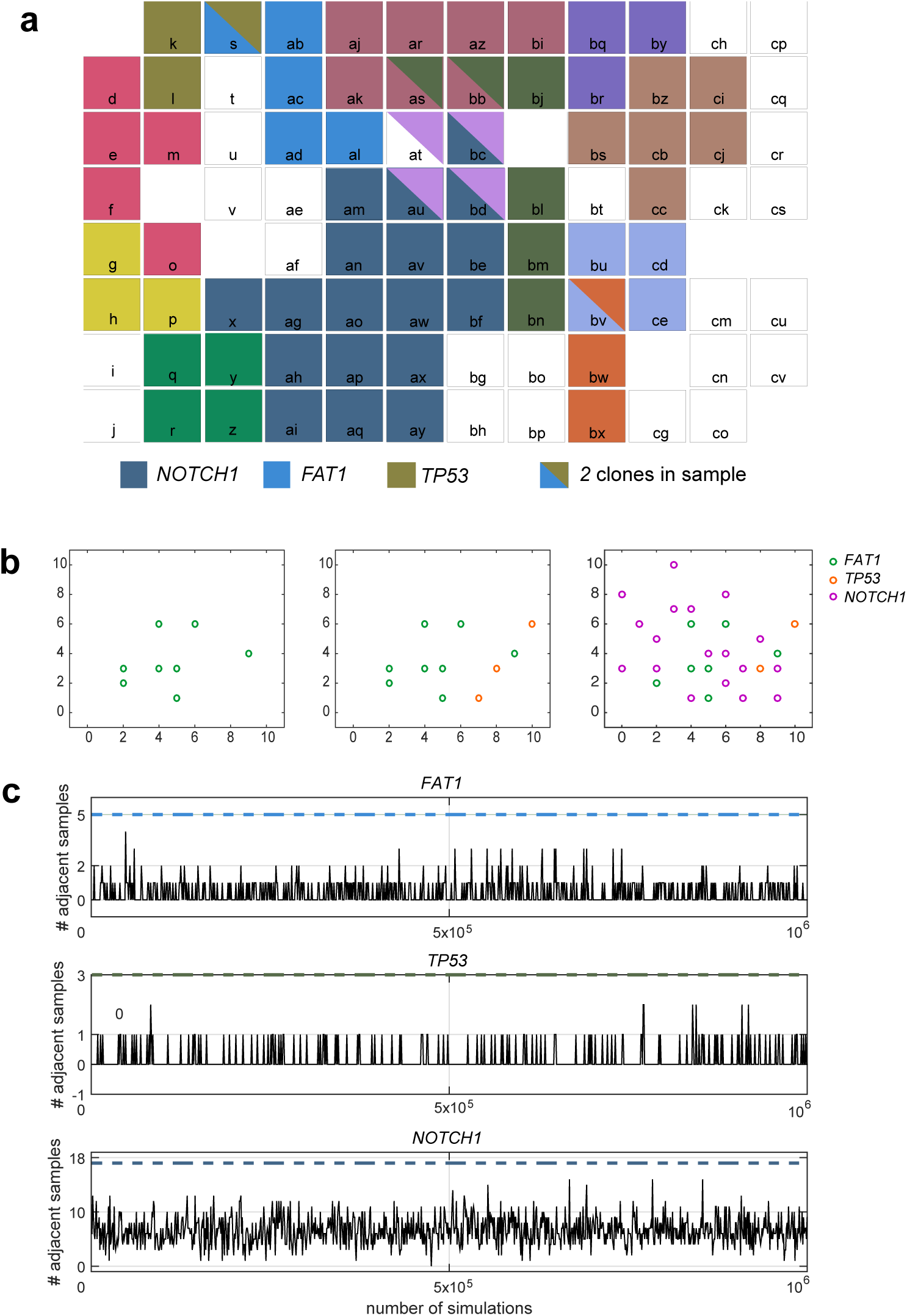
Size of largest selected mutant clones is consistent with competition. **a** Spatial map of large positively selected mutant clones in 0.05 mm^2^ samples from patient PD42776 (shown in **Figure 1)**. Clones spanning 3 or more adjacent samples are shown in different colors. The largest *NOTCH1*, *FAT1* and *TP53* mutant clones are indicated. **b** In silico permutation test of the likelihood that largest observed clone sizes arise by chance. In each simulation, the number of samples corresponding to the largest clone for each mutant is assigned to a spatial grid corresponding to that in each patient at random. In the example shown, 5 *FAT1* (green), 3 *TP53* (orange) and 18 *NOTCH1* (purple) mutations are randomly distributed across the grid. In this example the largest clusters spanned only 2 adjacent samples for *FAT1*, 0 for *TP53* and 3 for *NOTCH1*. **c** Result of 1 million simulations performed as shown in **b**. No simulations replicated all three of observed largest clone sizes, showing these are unlikely to have arisen by chance (p<1×10^−6^).

**Supplementary Figure 4:**
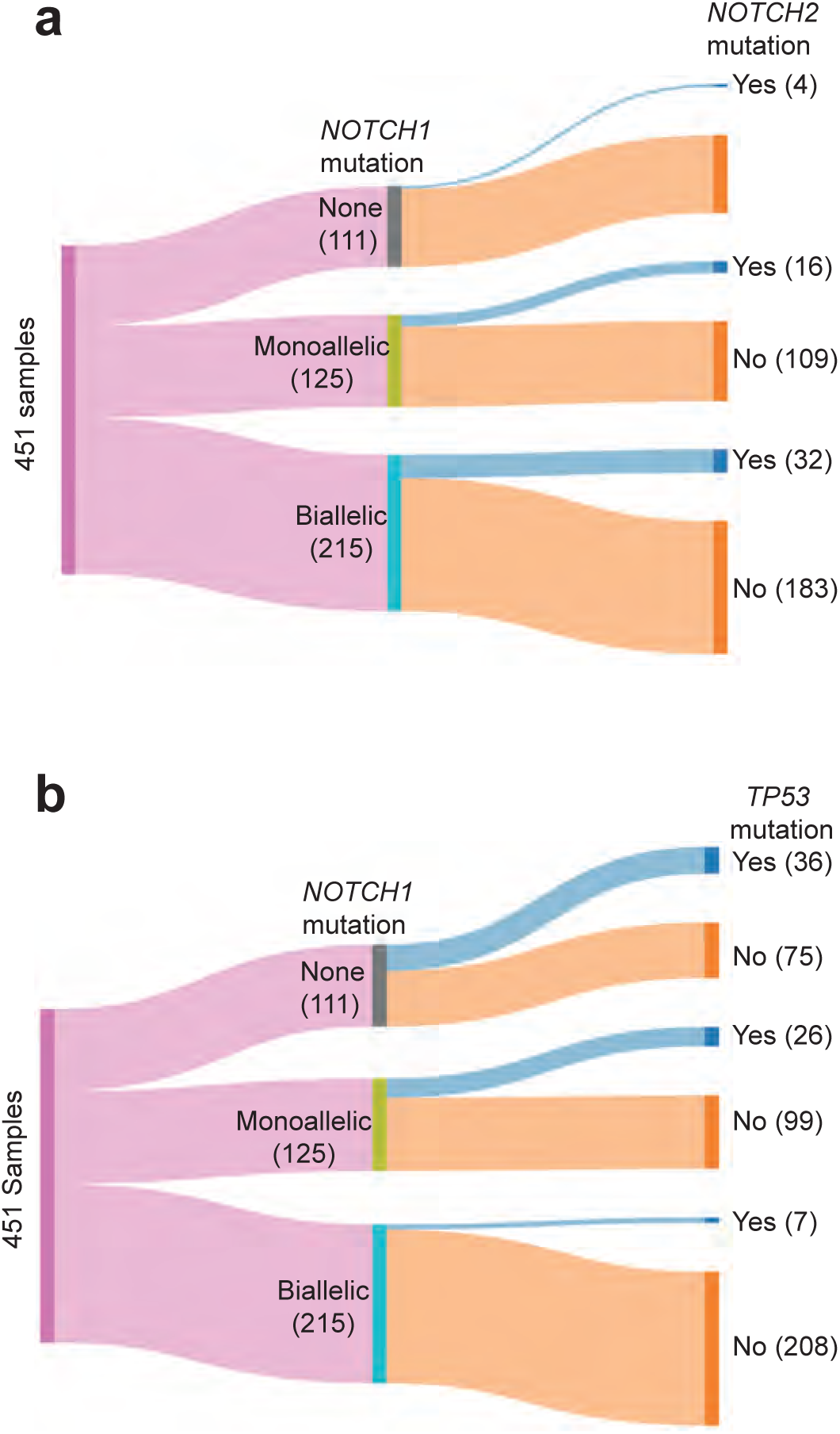
Numbers of *NOTCH2* and *TP53* mutations in *NOTCH1* mutant and wild type 0.05 mm^2^ samples. **a**, **b** Sankey plots showing numbers of samples which were NOTCH1 wild type or had mono– or bi-allelic *NOTCH1* mutation/LOH, that also carried *NOTCH2* (**a**) or *TP53* (**b**) mutations. There were significantly more *NOTCH2* mutants within samples carrying *NOTCH1* mutations compared to *NOTCH1* wild type samples (p=0.0026, Chi Square test). *TP53* mutations were enriched in *NOTCH1* wild type samples compared to those containing *NOTCH1* mutation/LOH mutants (p= 7.7×10^−9^, Chi Square test. Source data: Supplementary Table 7.

**Supplementary Figure 5.**
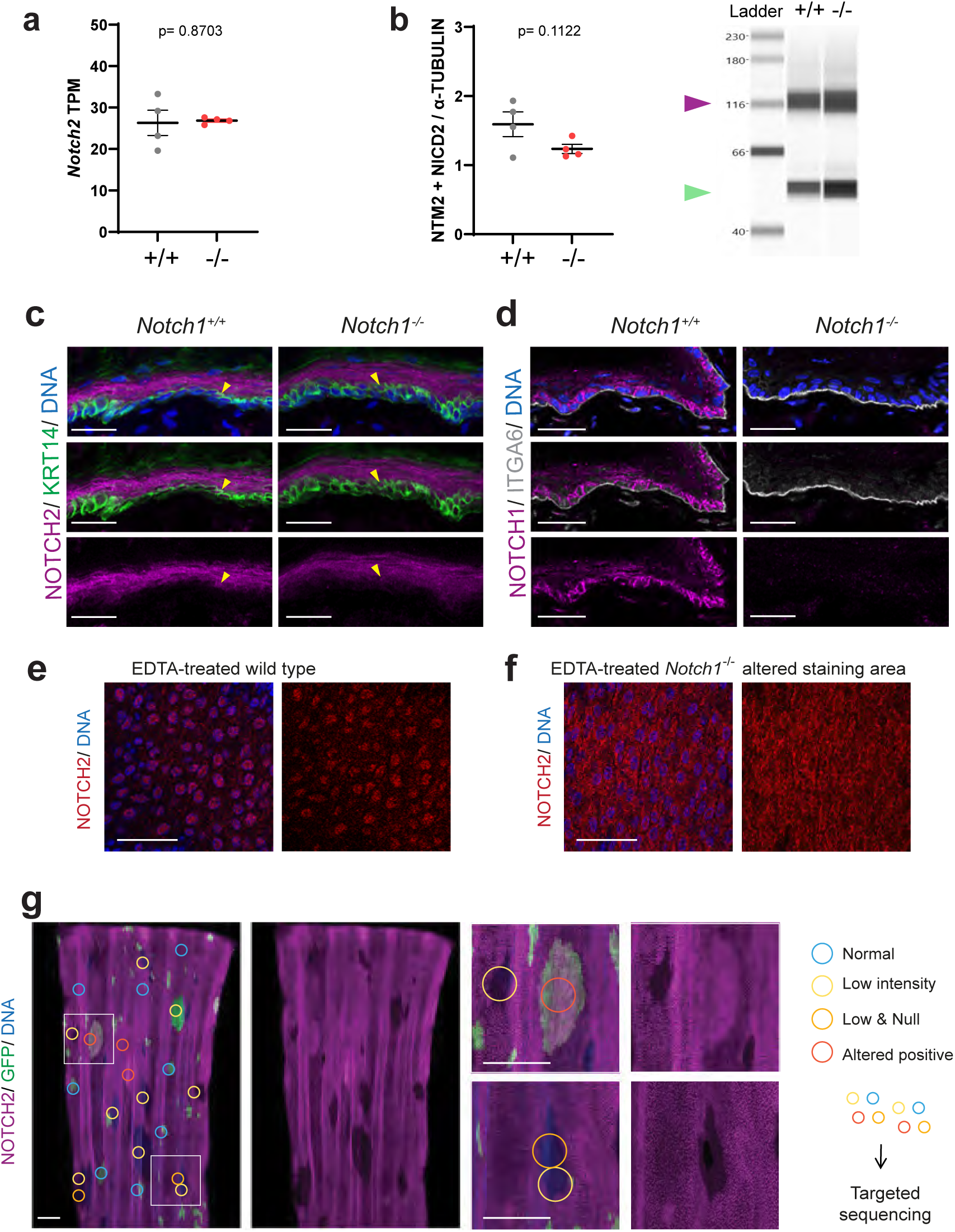
NOTCH2 expression identifies *Notch2* mutant clones. **a-d** Expression of Notch2 in wild type and Notch1−/−epithelium. **a** *Notch2* transcription in transcripts per million (TPM) from bulk RNAseq performed on fully induced *YFPCreNotch1^flox/flox^* (−/−) and uninduced littermate control mice (+/+) aged for 8 wk, mean ± SEM, each dot represents a mouse, n = 4 mice (Data from^7^). Two-tailed unpaired t test. Source data: Supplementary Table 20. **b** Immune Capillary Electrophoresis of NOTCH2 transmembrane/intracellular domain (NTM2 + NICD2, magenta arrow) and α-Tubulin protein (green arrow). Assay was performed on the same tissues as shown in **a**, mean ± SEM, each dot represents a mouse, n = 4 mice. Two-tailed unpaired t test. Source data: Supplementary Table 21.**c.** Cryosections of *Notch1^−/−^* and wild type control esophagus, stained for NOTCH2 (magenta), basal cell marker Keratin 14 (KRT14, green) and DNA (blue). Yellow arrow points at KRT14+ NOTCH2+ cells. Images are representative of three mice of each genotype. Scale bars, 30 μm. **d** Sections of *Notch1^−/−^* and wild type esophagus, stained for NOTCH1 (magenta), basal layer marker alpha-6 Integrin (ITGA6, grey) and DNA (blue). Images are representative of three mice of each genotype. Scale bars, 30 μm. **e, f** Rendered confocal z stacks of representative NOTCH2 control and altered positive areas in aged *Notch1^−/−^* esophagi treated with EDTA and stained for NOTCH2 (red) and 646 DNA (blue). **e**: Wild type NOTCH2 staining in *Notch2* wild type area, **f** altered NOTCH2 staining in *Notch2* mutant area. Top-down views showing first suprabasal cell layer. Scale bars, 50μm. **g** Whole mounts of highly induced *YFPCreNotch1^flox/flox^* mouse epithelium aner aging up to 78 weeks old and staining for NOTCH2 (magenta), GFP (green) and DNA (blue). Large ovoid areas of altered NOTCH2 staining (low intensity, low and null or altered positive staining classified as shown) were sampled for targeted sequencing (n=31 samples from 3 mice). White frames on the left images indicate magnified areas shown in the right-hand images. Scale bars, 500μm.

## Video Legends

**Video 1:** A spatial Moran simulation embodying experimental observations of biallelic *Notch2* mutant clones (black) in a heterozygous *Notch2* mutant background (green), time units are cellular generations. See **Supplementary Note** for model details and **Fig. 5** for stills of video.

**Video 2:** A spatial-Moran simulation of *Trp53* and *Notch1* single and double mutant clones evolving in a wildtype background (gray). *Notch1^−/−^*clones are black, *Trp53* mutant green, and *Trp53/Notch1* double mutants blue if *Trp53* mutation occurs first or yellow if *Notch1* mutation occurs first. T represents tissue generations. See **Supplementary Note** for model details and **Fig. 6** for stills of video.

## Methods

### Ethical Approval for animal experiments

Approval was given by the local ethical review committees at the Wellcome Sanger Institute and conducted according to Home Office project licenses PPL70/7543, P14FED054 and PF4639B40.

### Ethical approval for human study

The study protocol was ethically reviewed and approved by the UK National Research Ethics Service Committee East of England-Cambridge South, Research Ethics Committee protocol reference 15/EE/0152 NRES. Esophageal tissue was obtained from deceased organ donors. Written Informed consent was obtained from the donor’s next of kin.

### Human samples

A sample of mid-esophagus was removed within 60 minutes of circulatory arrest from 22 deceased organ donors at Addenbrooke’s Hospital (Cambridge, UK) and placed in University of Wisconsin (UW) organ preservation solution (Belzer UW® Cold Storage Solution, Bridge to Life, USA). Human esophagus epithelium was isolated and cut into a gridded array of 2 mm^2^ samples as described ^8^. DNA was extracted using QIAMP DNA microkit (Qiagen, 56304). Germline samples were collected from flash frozen esophageal muscle and DNA was extracted as for the epithelial samples.

DNA from a 2 mm^2^ sample was analyzed by nanorate sequencing (NanoSeq)^1^. As well as the 22 donors collected in this study, DNA from 9 donors reported in a previous study was also analyzed by NanoSeq.

### NanoSeq

NanoSeq is a method of single DNA molecule sequencing with an error rate less than five per billion base pairs, therefore providing an accurate estimation of genome-wide mutation burden^1^. Calling mutations from a single molecule of DNA also means that mutation burden estimates are not affected by the clonality of the sample. Nonetheless, to maximise consistency in sampling across donors, 2 mm^2^ samples were selected for NanoSeq if mutated for *NOTCH1* (as determined by targeted sequencing, see below), since this is the most common somatic cell genotype in aged esophageal epithelium. Substitution counts were corrected for genome-wide trinucleotide contexts and genome-wide burden estimated by dividing by the sequence footprint (**Supplementary Table 3**).

50ng of DNA from a 2 mm^2^ grid sample was used for library preparation according to^1^. Briefly, DNA was digested with mung bean nuclease, A-tailed, repaired and tagged. 0.3 fmol of indexed tagged library were sequenced with 14 PCR cycles before quantifying and sequencing on a NovaSeq6000 (Illumina) with 150 bp PE reads to give a median 0.9x duplex coverage. 30x coverage whole genome sequence of esophageal muscle from the same patient was used as the germline control for calling of SNPs and indels. Only samples with verifyBAMID alpha freemix <0.005 were used in downstream analysis. For both SNV and indels, only calls which passed all the defined filters were used https://github.com/cancerit/NanoSeq.

### Donor Genotyping

Donor genotypes were ascertained by whole-genome sequencing of donor muscle. A list of 34 ESCC risk alleles were obtained from the National Human Genome Research Institute-European Bioinformatics Institute GWAS (https://www.ebi.ac.uk/gwas/) and Open Targets (https://genetics.opentargets.org) databases. Associated genes are suggested for each single nucleotide polymorphism (SNP) using PheWAS and eQTL data from the Open Targets Genetics database. Risk allele frequencies for Great Britain (GBR), East Asia (EAS) and Africa (AFR) are quoted according to the 1000 Genomes Project Phase 3 to show donors in this study are representative of the UK population in terms of ESCC risk (**Supplementary Table 2**).

### 2 mm^2^ sample sequencing

For targeted sequencing of 2 mm^2^ grid samples 200 ng of genomic DNA was fragmented (average size distribution ∼ 150 bp, LE220, Covaris Inc), purified, libraries prepared (NEBNext Ultra II DNA Library prep Kit, New England Biolabs), and index tags applied (Sanger 168 tag set). Index tagged samples were amplified (6 cycles of PCR, KAPA HiFi kit, KAPA Biosystems), quantified (Accuclear dsDNA Quantitation Solution, Biotium), then pooled together in an equimolar fashion. 9 500 ng of pooled material was taken forward for hybridization, capture and enrichment (SureSelect Target enrichment system, Agilent technologies), normalised to ∼6 nM, and sequenced with 75 bp PE reads on HiSeq 2500. Samples were sequenced with one of two bait panels. The 74 gene bait panel covered the following genes: *ADAM29, ADAMTS18, AJUBA, AKT1, AKT2, APOB, ARID1A, ARID2, AURKA, BAI3, BRAF, CASP8, CCND1, CDH1, CDKN2A, CR2, CREBBP, CUL3, DICER1, EGFR, EPHA2, ERBB2, ERBB3, ERBB4, EZH2, FAT1, FAT4, FBXW7, FGFR1, FGFR2, FGFR3, FLG2, GRIN2A, GRM3, HRAS, IRF6, KCNH5, KEAP1, KMT2A, KMT2C, KMT2D, KRAS, MET, MUC17, NF1, NFE2L2, NOTCH1, NOTCH2, NOTCH3, NOTCH4, NRAS, NSD1, PCED1B, PIK3CA, PLCB1, PPP1R3A, PREX2, PTCH1, PTEN, PTPRT, RB1, RBM10, SALL1, SCN11A, SCN1A, SETD2, SMAD4, SMO, SOX2, SPHKAP, SUFU, TP53, TP63* and *TRIOBP.* The 32 gene bait panel covered a subset of these (*ADAM10, AJUBA, ARID1A, ARID2, CCND1, CUL3, DICER1, ERBB4, FAT1, FAT4, FBXW7, HRAS, KMT2C, KMT2D, KRAS, NF1, NFE2L2, NOTCH1, NOTCH2, NOTCH3, NRAS, NSD1, PIK3CA, PREX2, PTCH1, PTEN, RB1, SMAD4, SMO, TP53,* ZNF750).

BAM files were mapped to the human reference genome (GRCh37d5) using BWA-mem (version 0.7.17)^37^. Duplicate reads were marked using SAMtools (version 1.11)^38^. Depth of coverage (ca. 900x) was calculated using SAMtools to exclude reads which were: unmapped, not in the primary alignment, failing platform/vendor quality checks or were PCR/Optical duplicates. BEDTools (version 2.23.0) coverage program was then used to calculate the depth of coverage per base across samples^37^. BAM files of samples for nine published donors were then incorporated for all downstream analysis (**Supplementary Table 1**)^8^.

For targeted sequencing data, sub-clonal mutation variant calling was made using the deepSNV R package (also commonly referred to as ShearwaterML), version 1.26.1 commit d45c86c, available at https://github.com/gerstung-lab/deepSNV, used in conjunction with R version 3.5.2^8^. Variants were annotated using VAGrENT, https://github.com/cancerit/VAGrENT, version 3.3.3^39^.

Mutations detected using ShearwaterML were filtered by applying the following criteria:

○ Germline variants detected in the same individual were not considered called variants.
○ Adjustment for FDR and mutations demanded support from at least one read from both strands for the mutations identified.
○ Pairs of SNVs on adjacent nucleotides within the same sample are merged into a dinucleotide variant if at least 90% of the mapped DNA reads containing at least one of the SNV pair, contained both SNVs.
○ To prevent duplicate mutation counting Identical mutations found in multiple adjacent tissue biopsies were merged and treated as a single clone.

ShearwaterML was run with a normal panel coverage of approximately 42,000x for the 74-gene panel and around 12,000x for the 32-gene panel.

### 0.05 mm^2^ sample sequencing

0.05 mm^2^ samples were collected with circular biopsy punch (0.25 mm diameter, Stoelting Europe), in a rectangular array, with approximately a punch diameter between each sample. DNA was extracted using the Arcturus picopure kit (ThermoFisher).

Shearing and library prep was performed using the NEBNext®Ultra™ II Fragmentase System. DNA was hybridised using a bait panel described previously^40^ which contains many cancer driver genes. Material was subjected to 12 PCR cycles (Initial denaturation: 95C 5mins, 98C 30sec, 65C 30sec, 72C 2mins 12 cycles, 72C 10mins). Variants were called using the CaVEMan (version 1.15.1), http://cancerit.github.io/CaVEMan/, and cgpPindel (version 3.7.0), http://cancerit.github.io/cgpPindel, algorithms^41, 42^. For SNVs CaVEMan was run with the major copy number set to 10 and the minor copy number set to 2. Only SNVs which passed all CaVEMan filters were kept.

### Mutational signature analysis

Mutational spectra and signatures are described using the notation employed by the PCAWG Mutational Signatures working group^20^. COSMIC signature definitions (v3.2) (https://cancer.sanger.ac.uk/signatures/sbs/) were used for Single Base Substitutions (SBS), Double base Substitutions (DBS) and Indel signature classification. Using SigProfiler packages MatrixGenerator (v1.2.12), Extractor (v1.1.12), Assignment (v0.0.13), Plotting (1.2.2). The frequency of mutations within each trinucleotide context was calculated using SigProfiler within the SBS288^20^.

The maximum-likelihood implementation dNdScv (v0.0.1.0, https://github.com/im3sanger/dndscv) was used to identify genes under positive selection. dNdScv estimates the ratio of nonsynonymous to synonymous mutations across genes, adjusting for sequence composition and mutational spectrum. dN/dS values significantly greater than 1 indicate positive selection.

The proportion of mutant epithelium was calculated from mutant VAFs as described^8, 22^.

### Copy number calling

ASCAT (v3.2.0) was used to infer copy number from the 0.05 mm^2^ punch samples sequenced with a targeted bait panel. Allele counts in mutant genes under positive selection were generated for SNP positions from these normal samples allowing calling for both 0.05 mm^2^ and germline samples. These were used to derive LogR values and BAF values used to generate segment calls of allele-specific copy number. https://github.com/VanLoo-lab/ascat^43^.

### ‘Circle Plot’ visualisation of mutations in human esophagus

2mm^2^ samples were randomly selected to give a total sample area of 9cm^2^, for patients aged 30-59, 60-69 and 70-79. When two or more samples shared the same mutation at the same genomic coordinates, these mutations were collapsed into an individual event and their respective VAFs were merged.

The nesting of multiple mutations within samples was inferred from sequencing data when possible. For single samples containing a clonal mutation we used the pigeonhole principle to nest mutations^1^. If two or more mutations were shared by two or more samples the mutation they were considered nested if the VAF of one mutation was smaller than the VAF of other mutation(s) in all samples. If the VAFs of the shared mutations were not significantly different (tested using a Poisson test per sample) the samples were considered as mutants within the same clone, and nesting was assigned randomly. Finally, for multiple subclonal mutations within a single sample (e.g. two mutations with VAFs of 0.1 and 0.3), the presence of nesting was determined randomly, as long as the VAF of the largest mutation exceeded 0.05. Copy number information was not used to infer nesting.

To generate the ‘circle plot’ visualization, all clones were randomly distributed in space. Each circle is an observed non-synonymous positively selected mutation. Circle size scales to the corresponding variant allele frequency (VAF). The figure illustrates the number and density of mutant clones by age group.

### Mouse experiments

#### Animals and strains

Animals were 10 to 16 weeks old at the start of the experiments. Mice were housed in Specific and Opportunistic Pathogen Free unit with a 12-hour light/12-hour dark cycle. Lights were switched off at 19:30 and there was no twilight period. Ambient temperature was 21 ± 2 °C, and humidity 55 ± 10%. Mice were kept at 3–5 mice per cage, in cages measuring 365 mm × 207 mm × 140 mm (length × width × height), with a floor area, of 530 cm^2^. Individually ventilated caging (Tecniplast, Sealsafe 1284L) with 60 air changes/ hour was used. As well as Aspen bedding substrate, environmental enrichment with Nestlets, a cardboard tunnel and chew blocks was provided. Water and standard chow diet were provided ad libitum. Experiments were carried out with male and female animals, and no sex specific differences were observed. Animals were randomly assigned to experimental groups.

*YFPCreNotch1* triple mutant mice^7^ were generated by crossing B6.129X1-*Notch1^tm2Rko/GridJ^* mice purchased from the Jackson Laboratory with *Rosa26^floxedYFP^* and *AhCre^ERT^* mice ^27, 44, 45^. In these animals, *Cre* mediated recombination is induced by treatment with β-Naphthoflavone and Tamoxifen. An enhanced yellow fluorescent (EYFP) reporter is expressed from the Rosa 26 locus after *Cre* mediated excision of a *LoxP* flanked ‘stop’ cassette. Exon1 of *Notch1* is flanked by *LoxP* sequences so that *Cre* mediated recombination blocks *Notch1* expression.

*AhCre^ERT^Trp53^ffR245W-GF/wt^ Notch1^ffox/ffox^* conditional double mutant mice were generated by crossing *AhCre^ERT^Notch1^ffox/ffox^* mice with *Trp53^ffR245W-GFP^* animals^46^. In these animals, transient induction of *Cre* results in recombination at the *Trp53^ffR245W-GFP/wt^* locus with consequent expression of GFP, and at the Notch1 locus. Recombination efficiency differs between loci so clones lacking NOTCH1, expressing TRP53R245W and GFP, or both are generated.

#### Aging *Notch1* mutant mice

*YFPCreNotch1^flox/flox^* mice aged between 10-16 weeks-old were injected intraperitoneally (i.p.) with β-Naphthoflavone (MP Biomedicals, catalog no.156738) at 80 mg.kg^−1^ and tamoxifen (Sigma Aldrich, catalog no. N3633) at 0.25mg and collected at up to 78 weeks old. *Notch1^+/+^* or non-induced *YFPCreNotch1^flox/flox^* mice were used as wild type controls.

#### EdU Pulse Chase

EdU (5-ethynyl-2ʹ-deoxyuridine) incorporates during basal epithelial cell replication. 10 µg of EdU was injected intraperitoneally either 1 hour or 48h before tissue collection. Whole mount preparation and immunostaining were performed, including EdU detection using Click-iT EdU imaging kit (Life Technologies, catalog no. C10338 or C10340).

#### Immunostaining of esophageal sections

Mouse esophagus was flash frozen in tissue freezing medium (Leica catalog no. 14020108926). 10 µm thick cross-sections were fixed with 4% paraformaldehyde for 10 min, washed in PHEM and blocked in staining buffer (10% donkey serum, 0.5% BSA, 0.5% Triton X-100, 0.25% fish skin gelatin in PHEM). Sections were incubated overnight with the primary antibodies diluted in staining buffer (see **Supplementary Table 30** for antibody details). After PHEM washes, tissues were incubated for 3h with the secondary antibodies. Finally, after PHEM washes, tissues were incubated for an hour at room temperature with 1 μg.ml^−1^ DAPI to stain cell nuclei and mounted in Vectashield mounting media (Vector Laboratories, catalog no. H-1000).

#### Immune capillary electrophoresis

Protein extracts from fully induced esophageal epithelium and non-induced controls described in Abby et al., 2023 were used for NOTCH2 protein assay. Total protein was measured using Pierce™ BCA Protein Assay Kit (Thermo Fisher Scientific, catalog no. 10678484). Immune capillary electrophoresis was performed following manufacturer’s instructions for NOTCH2 and α-Tubulin detection using Wes Simple^TM^ (ProteinSimple). Quantification was performed using Compass for SW version 4.1.0. Anti-NOTCH2 targeting C terminus of the protein (Cell Signaling, catalog no. 5732) and anti-α-Tubulin (Cell signaling, catalog no. 2125) antibodies were used for this assay.

#### *Trp53, Notch1* double mutant mice

Both male and female adult mice of 10-16 weeks of age at the start of the experiments were induced by a single intraperitoneal (i.p) injection of 8 mgkg^−1^ ß-napthoflavone and 1.25 mg/ml tamoxifen and aged. Esophageal samples were collected at 4-, 12– and 24-wk time points.

#### Whole-mount preparation and immunostaining of mouse esophagus

Mouse esophagus was opened longitudinally before removing the underlying muscle layer with forceps. Esophageal tissue, comprising mucosa and submucosa, was incubated for 2h15 to 3 hours in 5 mM Ethylenediaminetetraacetic acid (EDTA) at 37 °C before peeling away the epithelium. The flattened epithelium was fixed in 4% paraformaldehyde for 60-75 minutes at room temperature under agitation then washed and stored in PBS at 4 °C ^4^. All subsequent steps were performed at room temperature. Tissues were incubated in staining buffer for a one-hour blockage (10% donkey serum, 0.5% bovine serum albumin, 0.5% Triton X-100, 0.25% fish skin gelatin in PHEM). This step was followed by overnight incubation with primary antibodies in staining buffer, three 30 min washes in 0.2% Tween-20 PHEM, and a 2-3 hour-long incubation with secondary antibodies in staining buffer (**Supplementary Table 25**). After washes in PHEM, tissues were incubated for 1h in 0.5 μM Sytox™ Blue nuclei counterstain (Biolegend, catalog no. 425305) or DAPI and mounted on slides using Vectashield media (Vector Laboratories, catalog no. H-1000).

#### Confocal microscopy

Immunofluorescence images were acquired using a Leica TCS SP8 confocal microscope with either ×10 or ×40 objectives. Images were acquired at a resolution of 1024 × 1024 pixels using optimal pinhole, line average 3-4 and a scan speed of 400 to 600 Hz. Image visualization and analysis were performed using Volocity 6.3 Image Analysis software (PerkinElmer).

#### Basal cell density

Wholemounted, stained tissues were imaged with a 40x objective using Leica TCD SP8 confocal microscope and basal cell density was analysed using Volocity 6.3 Image Analysis software (PerkinElmer). DAPI+ or Sytox™ Blue + nuclei were counted in the basal layer of NOTCH2 low intensity, null or control areas (**Supplementary Table 25**). Surface area measurements were performed using the Region of Interest tool.

#### Sequencing of aged esophageal epithelial wholemounts

Whole mounts from aged, induced *YFPCreNotch1^flox/flox^* or control mice were flattened and dissected into a grid of 2 mm^2^ contiguous biopsies. DNA extraction was performed using the QIAMP DNA microkit (Qiagen, catalog no. 56304) following the manufacturer’s instructions. Each sample was then subject to targeted sequencing.

#### Identification of *Notch2* mutant clones by immunostaining

We used a similar approach to that described for identifying *Notch1* mutant clones in a previous study^7^. Whole mounted epithelium was stained for NOTCH2, EYFP (using an anti-GFP antibody) and Sytox™ Blue or DAPI and imaged by confocal microscopy. Large, ovoid areas with either low intensity, increased intensity or null NOTCH2 staining suggested the presence of *Notch2* mutant clones. Similarly, large and ovoid areas with altered positive staining pattern such as predominantly cytoplasmic NOTCH2 protein also suggested the presence of *Notch2* mutations that would alter the cleavage or nuclear translocation of NICD2, as the EDTA incubation at 37 °C used to prepare the wholemounts, activates NOTCH2 cleavage^30^. Finally, YFP staining helped identifying putative mutant clones, appearing as large and ovoid areas devoid or fully stained with YFP. The mean intensity of fluorescence of NOTCH2 staining (red) and of DNA (blue) were measured on projected view of entirely imaged whole mounted epithelium (Leica TCS SP8 confocal microscope, ×10 objective). The ratio of mean intensity of red /mean intensity of blue (R/B ratio) of the whole tissue was used as reference to normalise the R/B ratio measured in each putative clones or control areas (**Supplementary Table 22**). Clonal surface area was measured using the region of interest tool.

#### Sequencing of *Notch2* mutant clones

Putative clones with altered NOTCH2 staining were sampled under a Fluorescent Stereo Microscope Leica M165 FC (Leica) using 0.25 mm diameter punch (Stoelting, catalog no. 57391). DNA extraction was performed using Arcturus® PicoPure® DNA extraction kit (Applied Biosystems, catalog no. 11815-00) following the manufacturer’s instructions. Targeted sequencing was then performed on each sample.

#### Targeted Sequencing of mouse samples

After multiplexing, samples were sequenced on Illumina HiSeq 2000 sequencer using paired-end 75-base pair (bp) reads. An Agilent SureSelect custom bait capture kit comprising 73 cancer related and Notch pathway related genes (baitset_3181471_Covered.bed, **Supplementary Table 14**) was used^7^.

For 2 mm^2^ samples the average depth of coverage across all genes was: *Notch1^+/+^*: 365x; *Notch1^−/−^*: 339x. For 0.05 mm^2^ samples the average depth of coverage across all genes was 228x.

Paired-end reads were aligned to the GRCm38 reference mouse genome with BWA-MEM (v.0.7.17, https://github.com/lh3/bwa)^37^ with optical and PCR duplicates marked using Biobambam2 (v.2.0.86, https://github.com/gt1/biobambam2).

Mutations in 0.05 mm^2^ samples from aged *YFPCreNotch1^−/−^* mice were analysed with CaVEMan (version 1.13.14) and with cgpPindel (version 3.3.0)^42^ for insertions and deletions. Sequence variants were annotated with VAGrENT (version 3.3.3)^39, 41^.

Mutations in 2 mm^2^ samples were also called with deepSNV (v1.26.1), https://github.com/gerstung-lab/deepSNV). The total coverage provided by the references used for the mouse bait kit was 22885x in a total of 34 germline samples.

In a gridded array of samples, some mutations can span between several adjacent biopsies. To prevent counting them multiple times and obtain accurate estimates of clone sizes and number, mutations extending over two or more biopsies were merged as individual calls^14,27^.

Selection was determined using dNdScv (v0.0.1.0) commit c3437e.

The proportion of cells carrying a mutation within each sample was calculated from the fraction of cells carrying a mutation within the sample, as described previously^8, 22^. The lower and upper bound estimates of the percentage of epithelium covered by clones carrying nonsynonymous mutations in each gene was then calculated. The proportion of mutant epithelium was derived from mean of summed VAF (capped at 1.0) of all samples in the esophagus.

## Statistical analysis

Data in graphs are shown as mean values ± standard error of the mean (SEM) unless otherwise stated. In general, P values <0.05 were considered statistically significant, but for dN/dS ratios calcuated by dNdScv, a global q<0.01 was considered statistically significant. All significance tests were 2 tailed. Each mouse experiment was performed using multiple biological replicates. The numbers of experimental replicates are given in figure legends and Supplementary Tables. Statistical tests described in figure legends were performed using GraphPad Prism software 8.3.1 and Python package SciPy 1.7.3 (https://scipy.org/citing-scipy/). No statistical method was used to predetermine sample size. Animals of the relevant genotype were allocated to experimental groups at random.

Two-dimensional permutation tests were performed to determine the likelihood that the observed spatial distribution of mutant clones in 0.05 mm^2^ samples of human esophagus arose by chance. From the spatial maps of each donor (Fig. 1b, Supplementary Fig. 2a, Supplementary Table 6), we determined the following:

i) The number of spatial arrayed samples sequenced in each donor
ii) The number of the largest clones carrying positively selected mutant genes (nM, usually 3).
iii) For the clones in ii) the size (sM, the number of adjacent samples harboring the same mutation)

The largest mutant clones were then modelled in silico. For nM clones, sM positions on the sample array were randomly and uniformly positioned on the 2D grid: the number of draws corresponded to the nM and the number of positions corresponding to sM for each mutant. The number adjacent positions occupied by the same mutant was determined to give the simulated cluster size, ssM, for each draw.

For each mutant M we then compared to the observed largest cluster size sM and simulated size ssM. If ssM > sM it is likely to that the observed clustering of the mutant occurred by chance. Conversely, if ssM < sM it is unlikely to the observed mutant clustering occurred by chance. An observed configuration for nM mutants is obtained by chance when SSM1 >SM1, SSM2>SM2, … SSMi >SMi in one simulation simultaneously for nM draws. 1 million simulations were performed for each donor. In no case was the observed size of the largest mutant clones generated, so p<10^−6^ that the observed largest clone sizes arose by chance for all donors.

## Data availability statement

The sequencing datasets generated during study are available at the European Genome-phemone Archive (EGA) with the following dataset accession numbers:

EGAD00001015256 – 2 mm^2^ samples, targeted

EGAD00001015258 – 0.25 mm^2^ samples targeted

EGAD00001015260 – Nanoseq

EGAD00001015261 – Nanoseq germline

Targeted sequencing of mouse tissue is deposited in the European Nucleotide Archive (ENA) under the following accession code:

ERP164438-Targeted sequencing of aged *Notch1^+/+^* mouse esophageal epithelium, targeted sequencing of aged *Notch1^−/−^* mouse esophageal epithelium, and targeted sequencing of aged *Notch1^−/−^* mouse esophageal epithelial microbiopsies.

## Supplementary Note

Here we discuss additional and modelling details relating to the mouse experiments described in the main text.

### 1. Spatial Moran-like models illustrate how epistasis shape the order of mutations

#### 3.1 Mutant Notch1/Notch2/Trp53 fitness interactions and clone fitness

The competition between clones in aging tissue and models of aging tissue have been extensively used in previous work^4^. Here we adapt those same approaches to explore mechanisms of gene interactions and compare them with qualitative outcomes from experimental studies presented here.

Briefly, the basic algorithm works as follows:

1. Loop through every cell in the grid in a randomized order. For each cell:
  a. Pick a neighbor of the cell
  b. Compare cell fitness
  c. A random number drawn from a uniform distribution from zero to one. If this is smaller than the difference of dividing and replacing cell fitnesses plus 0.5, the neighbor has their fitness, identity marker, and colour replaced by that of the dividing cell. For example, a cell with fitness of 0.5 attempting to replace a neighbour with a fitness of 0.5 will be successful 50 % of the time.
2. Go to step 1.

In the netlogo implementation ( https://ccl.northwestern.edu/netlogo/index.shtml), cells are modelled as patches on a square grid, and neighbours are considered to be four patches immediately adjacent to the dividing cell. Individual cells have the variables fitness (float), genotype (string), and color (used for visualisation). All cell fitnesses are initially set to 0, and genotype to wild type (“wt”). Mutations are introduced at an induction frequency specified by the user at each generation (‘netlogo tick’), by randomly altering cell fitness to a user set variable and genotype to a new string.

To explore epistasis, four additional genotypes are defined. In addition to “wt”, “a”, “b”, “ab”, and “ba” represent two mutations that can occur in a single cell, in two different orders. The specific mutation that occurs is determined randomly, and each cell can acquire a single mutation no more than once. The fitness values of “a” and “b” are user specified (F_a_ and F_b_ respectively), whilst both “ab” and “ba” are the sum of individual fitnesses and an additional value representing epistatic interaction (F_e_) i.e. F_ab_ = F_ba_ = F_a_ + F_b_ + F_e_

Individual parameters are specified manually to illustrate fundamental principles of gene interaction and not fit to experimental data. The probability of a cell acquiring a mutation in a single generation as 1 in 100,000. We seek explicitly to qualitatively demonstrate the mechanisms of gene interaction with these models.

In previously simulated environments with no epistatic interactions (F_e_ = 0), clones bearing a and b mutation initially takeover the tissue, before being colonised and replaced by clones bearing both mutations with different orders in similar proportions. Once takeover is complete, cells compete neutrally, and overall fractional takeover remains steady.

To explore apparent epistatic interactions between *Notch1* and *Notch2* and *Notch1* and *Trp53* we specified different values to represent the respective independent fitness effect and the epistatic contribution to fitness. *Notch2* mutants fail to persist in a wildtype background but colonise the esophagus in a *Notch1^−/−^* background. To explore, this we specify that mutant *Notch1* and *Notch2* have independent fitnesses of 0.1 (positive selection) and 0 (neutral) respectively (F_NOTCH1_ = 0.1, F_NOTCH2_ = 0), and the epistatic contribution is 0.1 (F_e_ = 0.1). As expected, *Notch2* mutant clones fail to persist and expand in a wildtype background, but expand readily in within *Notch1* mutant clones, eventually taking over the whole tissue. The epistatic interaction effectively enforces one order of mutations, as *Notch2* mutants fail to expand and so are unlikely to acquire a second mutation. Almost all final genotypes represent *Notch1* mutation followed *Notch2* mutation.

Mutant *Trp53* and *Notch1* show patterns consistent with genetic dominance of the fitness advantage of *Notch1*. However, both mutant genes individually colonise the tissue, with *Notch1* mutant clones more common than clones bearing *Trp53* mutation. To model this we specify mutant *Notch1* and *Trp53* independent fitnesses as 0.1 and 0.07 (F_NOTCH1_ = 0.1, F_TP53_ = 0.07), and a negative epistatic contribution of –0.07 (F_e_ = –0.07). This limits the overall fitness of double mutant clones to the higher fitness of single mutant *Notch1*.

This has several impacts on the evolving system. Initially single mutant clones expand in the tissue over time. Eventually, these single mutant clones collide, at which point mutant *Notch1* clones outcompete mutant *Trp53* clones causing them to regress, and the overall proportion of mutant *Trp53* tissue to drop, consistent with observation. Double mutations arise in the tissue, but their expansion depends on the order of mutation. *Trp53* mutations in a *Notch1* mutant background fail to expand as they have no advantage over adjacent *Notch1* cells and can only persist if they occur at the growing edge of a clone where they can successfully expand into wild type or single *Trp53* mutant background. In contrast, *Notch1* mutation allows clonal expansion in a *Trp53* background due to the increase of fitness relative to surrounding parental cells. This leads to a small fraction of *Trp53*/*Notch1* clones when the tissue is completely colonized.

These findings are consistent with observed human sequencing data where rare *NOTCH1*/*TP53* double mutant clones can be found. It further illustrates an important concept in clonal evolution. Though both double mutants are equally fit, only the double mutant clones where *TP53* mutation has occurred first can successfully expand and persist. This reflects the need for each individual mutational event to increase fitness to drive clonal expansion. This concept also holds for *NOTCH1*/*NOTCH2* mutant clones, where both orders of mutation are ultimately equally fit, but only *NOTCH2* mutations that are preceded by a *NOTCH1* mutation successfully colonize the tissue. Here the neutrality of the *NOTCH2* single mutant prevents persistence in the tissue, effectively acting as a barrier to the creation of the double mutant. This illustrates the need for a “fitness escalator” for clonal expansion where each subsequent mutation must offer a relative fitness advantage for combined mutations to spread.

#### 1.2 Mechanisms of mutant *Notch2* expansion through ‘defensive fitness’

Our model of clonal expansion uses a single parameter to determine selective advantage, fitness. Fitness enables clonal expansion through competition with less fit neighbours but does not alter proliferation or stratification between cells of equal fitness. Once the tissue is colonised by clones of equal fitness competition becomes neutral and proliferation and stratification. Experimentally we observe that stratification in *Notch2* double mutants (expressing no NOTCH2 protein) is reduced, inconsistent with this model. Fitness is used in two different checks in our code, determining whether a cell stratifies and whether a cell divides. To explore whether we could explain mutations that increase fitness and alter stratification or proliferation, we separated fitness into two independent variables, termed F_offensive_ and F_defensive_, to represent the parameters used for proliferation or stratification respectively.

Increasing either of these parameters enables clonal expansion in our model system, albeit at a slower rate than when both are increased equally. We find however that increasing F_defensive_ in isolation reduces stratification within mutant clones as neighbouring cells of equal fitness remain less able to stratify. Similarly increasing F_offensive_ in isolation increases proliferation within mutant clones. This suggests a simple mechanism of *Notch2* mutant clonal expansion where loss of *Notch2* enables clonal expansion by preventing stratification specifically.

